# A GGDEF domain serves as a spatial on-switch for a phosphodiesterase by direct interaction with a polar landmark protein

**DOI:** 10.1101/2021.08.12.456111

**Authors:** Tim Rick, Vanessa Kreiling, Alexander Höing, Svenja Fiedler, Timo Glatter, Wieland Steinchen, Georg Hochberg, Heike Bähre, Roland Seifert, Gert Bange, Shirley K. Knauer, Peter L. Graumann, Kai M. Thormann

## Abstract

In bacteria, the monopolar localization of enzymes and protein complexes can result in a bi-modal distribution of enzyme activity between the dividing cells and heterogeneity of cellular behaviors. In *Shewanella putrefaciens*, the multidomain hybrid diguanylate cyclase/phosphodiesterase PdeB, which degrades the secondary messenger c-di-GMP, is located at the flagellated cell pole. Here we show how PdeB polar recruitment is mediated by direct interaction between the inactive diguanylate cyclase (GGDEF) domain of PdeB and the C-terminal FimV domain of the polar landmark protein HubP. We demonstrate that this interaction is crucial for full function of PdeB as a phosphodiesterase. Thus, the GGDEF domain serves as a spatially controlled on-switch that effectively restricts PdeBs activity to the flagellated cell pole. We further show that PdeB regulates abundance and activity of at least two crucial surface-interaction factors, the BpfA surface adhesion protein and the MSHA type IV pilus. The heterogeneity in c-di-GMP concentrations that is generated by differences in abundance and temporal polar appearance of PdeB as well as by bi-modal distribution after cell fission orchestrates the population behavior with respect to cell-surface interaction and environmental spreading.

**Significance:** Phenotypic heterogeneity benefits the proliferation of microbial populations in changing environments. Such heterogeneity can be created by recruitment of enzymatic activity to specific cellular compartments, e.g., the cell pole. Here we show how a GGDEF domain of a multidomain phosphodiesterase has adopted the function as a spatial on-switch that is specifically activated upon direct interaction with a polar landmark protein.

## Introduction

Cyclic mono- or oligonucleotide second messenger molecules are ubiquitously occurring in cells from all three domains of life and are implicated in a wide array of signaling networks at the single or multiple cell level. The second messenger cyclic bis-(3′–5′)-cyclic diguanylic acid (c‑di‑GMP) has emerged as an important signaling molecule in a multitude of bacterial species (1–3). c-di-GMP is synthesized from two molecules of GTP by specific diguanylate cyclases (DGCs), which are characterized by a signature GGDEF pentapeptide motif within their reactive center. Degradation via hydrolysis of c-di-GMP is mediated by phosphodiesterases (PDEs) that contain either signature EAL or HD-GYP domains and convert c-di-GMP to linear pGpG or to GMP, respectively (reviewed in (4)). These domains may occur as stand-alone proteins, however, more frequently they are found as parts of larger multidomain proteins together with various sensor input or output domains. Often, both GGDEF and EAL domains jointly occur as part of multidomain (sensor)proteins, which may localize in the cytoplasm or may be membrane-bound. The processes c-di-GMP-mediated regulation is involved in are highly diverse. In numerous bacterial species this secondary messenger plays a key role in controlling the shift between sessile and planktonic lifestyle. In addition, c-di-GMP is implicated in regulation of the cell cycle, cellular development, differentiation, and virulence (reviewed in (1, 5)).

Mechanisms that ensure appropriate and specific phenotypic responses to c-di-GMP include timing of PDE/DGC or effector production, effector affinity to c-di-GMP, or spatial control by sequestration of the regulator/effector components into functional clusters or to specific cellular compartments (3, 6, 7). In several bacteria, DGCs and/or PDEs are specifically recruited to the cell poles. In *Caulobacter crescentus*, which has an asymmetric cell cycle in which a sessile stalked cell produces a motile swarmer cell, c-di-GMP is an important regulatory factor of cell cycle control and pole morphogenesis (8–10). One key factor is the DGC PleD, which is localized to and activated at the pole of the sessile stalked mother cell. In contrast, an active PDE PdeA is present at the opposite pole of the newborn swarmer cells where it keeps the c-di-GMP concentrations low (10), leading to a bimodal distribution between mother and daughter cell upon division (11). A similar bimodal distribution of c-di-GMP was described in *Pseudomonas aeruginosa* (12), where the PDE Pch is recruited to and activated at the flagellated cell pole via phosphorylation by the chemotaxis histidine kinase CheA. Accordingly, the Pch-induced asymmetry results in low c-di-GMP concentrations of the already flagellated cell promoting active swimming (12) or detachment of surface-associated cells, leaving the still attached offspring behind (13). This process, termed ‘Touch-Seed-and-Go’, enhances spreading of the population and demonstrates how heterogeneity in c-di-GMP concentrations may benefit proliferation of the species.

In *Shewanella oneidensis* and *S. putrefaciens*, the phosphodiesterase PdeB is a main regulator of the transition from planktonic to sessile life style in response to a yet unknown environmental signal (14, 15). PdeB consists of a putative periplasmic signal receptor domain, flanked by two transmembrane regions, and cytoplasmic section with a HAMP and a PAS domain followed by a GGDEF and an EAL domain (see **Fig. 1a**). The EAL domain is crucial for overall function of PdeB, however, full activity *in vivo* requires every single domain of the protein (15). *Sp*PdeB is directly recruited to the flagellated cell pole by the polar landmark protein HubP. HubP/FimV orthologs occur in a number of different bacterial species (e.g., *P. aeruginosa*, *Vibrio* and *Shewanella*). In these species these proteins coordinate a range of different cellular processes involving, but not restricted to, flagellation, swimming motility, and chemotaxis (16–23), type IV pili assembly and activity (24–29), and recruitment of the origin of replication to the cell pole (15, 16). The polar localization of PdeB solely depends on its GGDEF (DGC) domain, which directly interacts with the C-terminal FimV domain of HubP (15). Assembly of a novel PdeB cluster at the pole of the non-flagellated daughter cell normally occurs long after cell fission, suggesting that PdeB induces an asymmetry with respect to c-di-GMP concentrations between the daughter cells in dependence of environmental signals.

**Figure 1:**
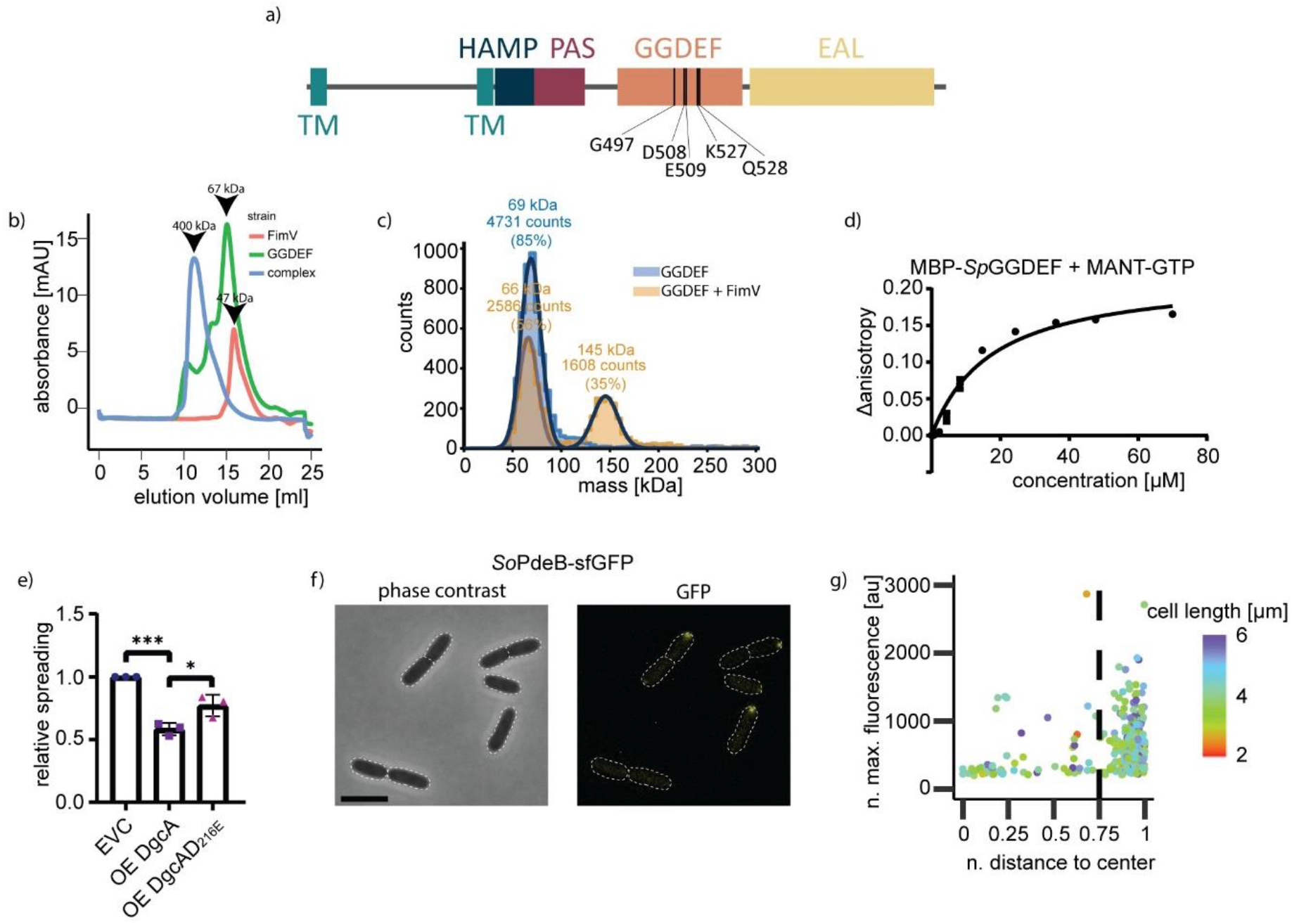
Common features of PdeB. **a)** Domain organization of PdeB. The position of important residues within the DGC/GGDEF domain are marked. **b)** SEC interaction studies of MBP-GGDEF_*So*PdeB_ with FimV_*So*HubP_. Both proteins form a complex *in vitro*. **c)** iSCMAS analysis to determine the stochiometry of the MBP-GGDEF_*So*PdeB_/FimV_HubP_ complex. While under high protein concentrations a large complex is present (b), iSCAM assays indicates a complex of 2 FimV to 2 GGDEF. **d)** Binding of MANT-GTP to the GGDEF domain of *So*PdeB by fluorescence anisotropy measurements. **e)** Residue D216 of DgcA is crucial for DGC activity, as shown by soft agar motility assay. This residue usually forms a salt bridge with a conserved C-terminal lysine and all PdeB homologues encode glutamic acid instead of aspartic acid at this position. **f-g)** Localization behavior of *S. oneidensis* PdeB-sfGFP was tested by fluorescence microscopy and is shown as micrographs (scale bar = 5 μm) and scatter plots.

Here, we characterized the interaction of PdeB with HubP between the two proteins and mapped the interaction surface between the GGDEF domain of PdeB and FimV of HubP. *In vivo* and *in vitro* analyses strongly indicate that the interaction with HubP at the cell pole is crucial for PdeB function, which therefore is only fully active upon recruitment to the cell pole. This and the difference in abundance of PdeB enhances the heterogeneity of the c-di-GMP concentrations within the population, promoting a diversity of cellular c-di-GMP-dependent processes.

## Results

### Mapping the FimV_HubP_/GGDEF_PdeB_ interaction site

Our previous studies showed that in *S. putrefaciens* the GGDEF domain of PdeB (GGDEF_PdeB_) directly interacts with the C-terminal FimV domain of the polar landmark HubP (FimV_HubP_) to mediate polar localization of PdeB (15). However, it remained unclear how the two domains interact and if this has functional consequences for the activity of PdeB. Therefore, we set out to identify the critical residues mediating the interaction between GGDEF_PdeB_ and HubP_FimV_. For this, we used a two-pronged complementary approach: As we showed that other *S. putrefaciens* GGDEF-domain proteins do not interact with FimV, we reasoned that GGDEF_PdeB_ has distinctive features not present in canonical GGDEF domains and may therefore be involved in mediating the interaction with FimV_HubP_. In parallel, we directly mapped the interaction surface by *in vitro* protein studies.

### Identification of common features in *Shewanella* PdeB

Potential orthologs to PdeB are present in numerous different species of *Shewanella*. To first determine whether polar recruitment may be a common feature of *Shewanella* PdeB, we localized PdeB from *S. oneidensis* (*So*PdeB (14)) by chromosomally fusing *pdeB* to *sfgfp*, leading to production of stable fluorescently labeled *So*PdeB-sfGFP (**Fig. S1d**). As in *S. putrefaciens*, *So*PdeB-sfGFP exclusively localized to the cell pole (**Fig. 1f, g**). To investigate whether *So*PdeB also interacts with FimV_HubP_, we heterologously produced and purified the GGDEF domain of *So*PdeB and found that it forms a complex with both the FimV domains of *S. oneidensis* (*So*FimV) and *S. putrefaciens* (*Sp*FimV) after size exclusion chromatography (**Fig. 1b**). According to interferometric scattering mass spectrometry (ISCAMS) analysis the complex had a molecular mass of about 145 kDa (**Fig. 1c**), which would fit to a 2:2 stoichiometry. We therefore hypothesized that PdeB/HubP interaction via the GGDEF and FimV domains is a common feature in *Shewanella*, indicating that the residues within the FimV-GGDEF interaction surface are conserved between *S. oneidensis* and *S. putrefaciens* and maybe further *Shewanella* species. Therefore, the sequences of 50 putative PdeB orthologs from various species of the genus *Shewanella* were used to create a position-based weight map for PdeB, which was then compared to that of *bona fide* cyclases (30). The critical residues required for GTP binding and stabilization of the transition state are present in all PdeB homologs, however, the c-di-GMP-binding inhibitory site (I-site) is degenerated (GxxE in PdeB instead of RxxD in canonical GGDEF domains) and the R” region at the N-terminal region is fully missing in GGDEF_PdeB_ (**see Fig. S1a**). Accordingly, fluorescence polarization assays showed that purified the purified GGDEF domain of *S. putrefaciens* was able to bind GTP at a KD value of approximately 16.77 ± 4.33 μM, but not c-di-GMP (**Figs. 1d; S1b, c**). To determine whether GTP binding has a role in PdeB activity, we mutated chromosomal *pdeB* in its c-di-GMP-binding site (PdeB_GGAAF_). The c-di-GMP concentrations of the mutant bearing the substitution were significantly higher than those of wild-type cells (**Fig. S1e**), indicating that GTP binding to the GGDEF domain positively regulates PdeB activity.

As a second unusual feature, GGDEF_PdeB_ possesses a glutamic acid residue at position a position where canonical GGDEF domains have a conserved aspartic acid (E467 in *S. putrefaciens* PdeB; *Sp*PdeB). This variation is conserved among all PdeB homologues and may induce structural differences since the conserved aspartic acid usually interacts with a lysine (K578 in *Sp*PdeB) at the far C-terminus of the GGDEF domain. To test if the E467 variation represses DGC activity, we introduced a corresponding D to E substitution into the DGC VdcA from *Vibrio cholerae* (VdcA_D216E_), which has previously been shown to be active in *Shewanella* (31). VdcA or VdcAD216E were ectopically produced in *S. putrefaciens*, and a decrease in flagella-mediated spreading through soft agar was scored as a measure for DGC activity. While production of wild-type VdcA limits flagella-mediated spreading of *S. putrefaciens* through soft agar as expected, the overproduction of the DgcA_D216E_ variant had a significantly smaller effect (**Fig. S1d**). We thus concluded that the E467 variation negatively affects the enzymatic inactivity of the GGDEF domain from PdeB and, thus, may account for the inactivity of GGDEF_PdeB_ as DGC *in vivo* and *in vitro* (14, 15).

### Identification and prediction of the FimV_HubP_/GGDEF_PdeB_ interaction site

To determine the protein interaction surface between GGDEF_PdeB_ and FimV_HubP_, the GGDEF/FimV complex from *S. putrefaciens* was purified and analyzed by hydrogen–deuterium exchange mass spectroscopy (HDX-MS) (**Fig. 2a**). The assay revealed that residues 1047–1107 of HubP exhibited reduced deuterium content in the presence of PdeB_GGDEF_, indicating that this region serves as the docking site for PdeB. However, no HDX result could be obtained for PdeB_GGDEF_. As a complementary approach, we therefore applied cross link-mass spectroscopy (CL-MS) to analyze the protein complex of FimV_HubP_/GGDEF_PdeB_. This assay was conducted using *S. oneidensis* FimV_HubP_/GGDEF_PdeB_, which could be produced and purified at higher yields. We found that a peptide encompassing the residues 1053 – 1088 of *So*HubP was chemically cross-linked to the peptide encompassing the residues 566 – 577 of *So*PdeB (**Fig. 2b; S2a**). The peptide identified by CL-MS in FimV_HubP_ in *S. oneidensis* matched the region identified by HDX in *S. putrefaciens* FimV_HubP_-GGDEF_PdeB_ (**Fig. 2b**).

**Figure 2:**
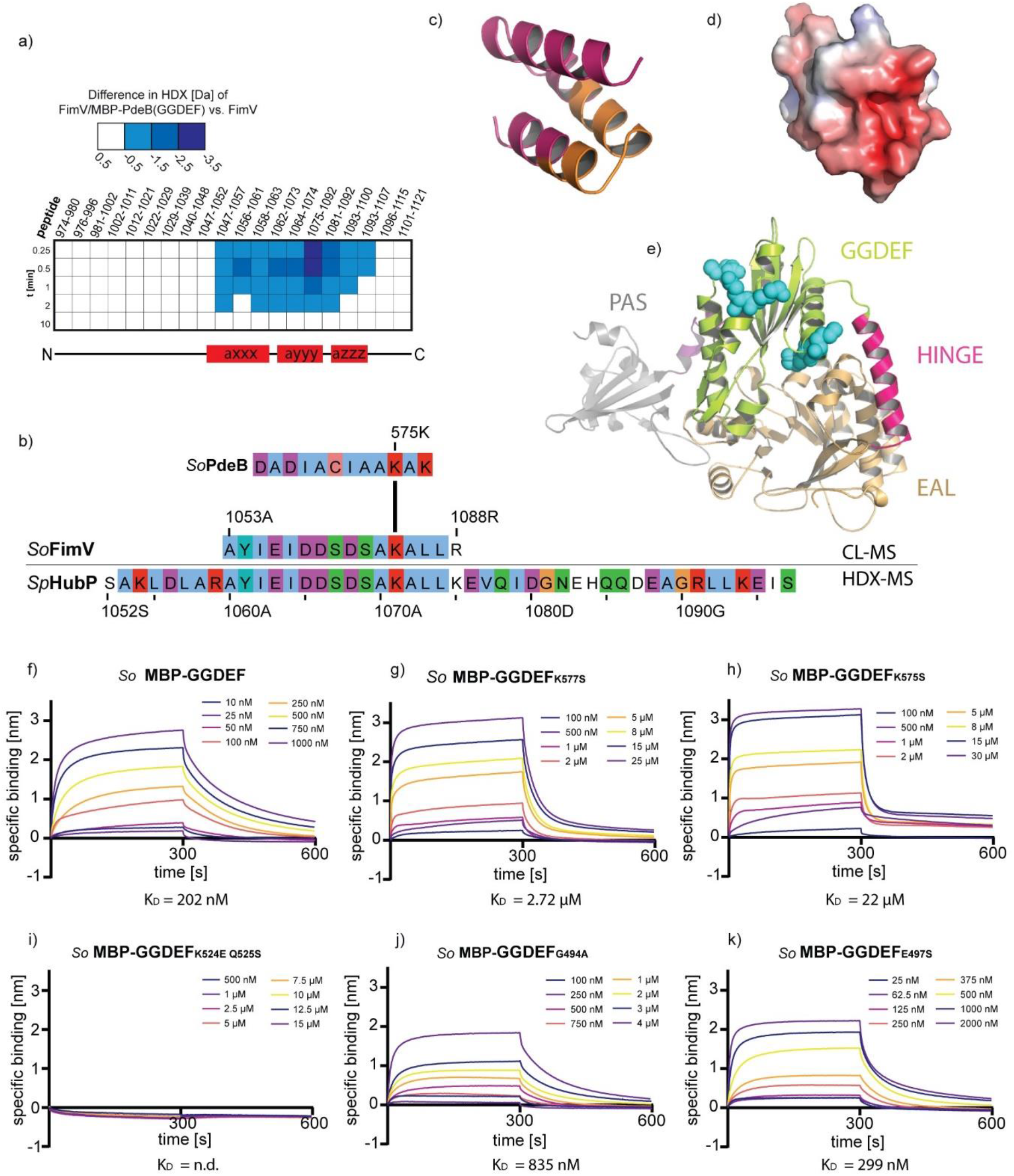
Interaction surface of PdeB and HubP. **a-b)** Regions that are involved in the interaction of PdeB and HubP were identified using HDX-MS (a) and CL-MS (b). The sequences of *Sp*HubP and *So*HubP were aligned using clustal omega and JalView. **c)** The structure of the C-terminal domain of *Sp*HubP was predicted using the swiss-model algorithm and the peptide found by CL-MS is colored in orange. **d)** The electrostatic surface potential was calculated using the APBS electrostatic tool and pymol. Red coloring indicates a negative surface charge. **e)** The structure of the PAS-GGDEF-EAL region of PdeB was predicted using Phyre2. The N-terminal PAS domain is colored in gray, the GGDEF domain in green and the C-terminal EAL domain in light orange. Residues within the GGDEF domain that we identified to be involved in the PdeB-HubP interaction are colored in cyan. **f)** The dissociation constant for the *So*PdeB-*So*HubP interaction was determined using BLI assays. **g-k)** Residues that are involved in respective interaction were substituted as indicated and the K_D_ value was determined using BLI assays. All mutations decreased the affinity of *So*GGDEF to *So*FimV.

The data obtained by the HDX- and CL-MS approaches allowed a definition of the interaction surface between both proteins. The structure of *Shewanella* FimV_HubP_, predicted based on that of the highly homologous FimVc-domain from *Pseudomonas aeruginosa* (26), showed a distinct negative surface charge of the FimV domain at the putative docking site of GGDEF_PdeB_ (**Fig. 2c,d**). We therefore assumed that the interaction surface of the domain includes critical positively charged residues. Correspondingly, the GGDEF_PdeB_ peptide that was identified by CL-MS to interact with FimV_HubP_ harbors two lysines, K575 and K577 (**see Fig. S1a**), which are fully conserved in all *Shewanella* PdeB proteins. K577_PdeB_ is variable in canonical GGDEF-domains while K575_PdeB_ is a conserved lysine that interacts with a conserved aspartic acid at the N-terminus of the GGDEF motif and is therefore functionally relevant (30). To identify additional residues potentially involved in polar recruitment of PdeB, we used the position-based weight map of PdeB homologs to screen for further surface-exposed, positively charged amino acids, which are not present in other, canonical GGDEF domains. According to these assumptions, the residue pair K524_PdeB_ and Q525_PdeB_ turned out as potential candidate, which in canonical GGDEF domains consists of a negatively charged glutamic acid followed by an arginine. While K524_PdeB_ is fully conserved in all PdeB homologues, the following residue Q525_PdeB_ is variable but always a large hydrophilic amino acid that is not negatively charged (**Fig. S1a**). As, according to structural predictions, both residues are near the degenerated inhibitory site (I-site; **Fig. 2e**) of GGDEF_PdeB_, we further included residues within this site to map the FimV/GGDEF interaction surface. Thus, the HDX-, cross-linking and *in silico* approaches identified several amino acid residues in GGDEF_PdeB_ that are putatively involved in mediating the interaction with HubP_FimV_.

### The far C-terminal region and degenerated I-site of GGDEF_PdeB_ mediate interaction with HubP_FimV_c

As the next step, we wished to determine whether the prediction of the GGDEF_PdeB_/FimV_HubP_ interaction surface was correct and performed *in vitro* interaction studies. Due to the better purification properties of *S. oneidensis* GGDEF_*PdeB*_ and the conserved interaction between GGDEF_PdeB_ and FimV_HubP_ from *S. putrefaciens* and *S. oneidensis*, the *in vivo* studies were conducted using *S. oneidensis* GGDEF_PdeB_. To this end, mutant variants of GGDEF_PdeB_ bearing amino-acid substitutions at the desired position were purified. To investigate if K575 and K577 in GGDEF_PdeB_ are involved in mediating interaction to FimV_HubP_, we used variants in which the appropriate lysine residues were substituted with a serine (K575S and K577S). A potential effect of the residues within or close to the degenerated I-site of GGDEF_PdeB_ was determined by introducing a series of single substitutions (G494A; E497S; Q525S; K524S) and a K524E Q525S double substitution into the protein. To verify structural integrity and functionality of the mutated proteins we showed that all *So*PdeB_GGDEF_ variants elute at the same volume after size exclusion chromatography (SEC) and still bind MANT-GTP with roughly the same affinity as wild-type *So*PdeB_GGDEF_ (**Figs. S3a-i; Fig. S2c**). Interaction of the wild-type and mutant proteins to *So*FimV_HubP_ was then determined by bio-layer interferometry (BLI) (**Figs. 2f–k; Fig. S2d,e**). Wild-type GGDEF_PdeB_ bound to FimV_HubP_ with high affinity at a K_D_ value of around 200 nM (**Fig. 2f**). Compared to the wild-type version, both substitutions in the conserved lysins strongly decreased the affinity to FimVc and resulted in a 100-fold (K575S) and a 10-fold (K577S) increased K_D_-value (**Fig. 2g,h**). Also all substitutions targeting the region around the I-site negatively affected the affinity of the protein variant towards FimV to various extends (**Fig. 2i–k; Fig. S2d,e**). The *So*GGDEF_K527E-Q528S_ double substitution even completely suppressed FimV_HubP_-GGDEF_PdeB_ interaction *in vitro* (**Fig. 2i**). Determination of the K_D_ value for the FimV-GGDEFK524S variant by BLI assays was not possible due to slight aggregation of the latter protein (**Fig. S2f**). However, pull-down assays suggest that also this variant does not effectively bind FimV (**Fig. S2g**). Taken together, HDX, cross-linking and mutational analysis allowed us to predict the interaction surface between GGDEF_PdeB_ and FimV_HubP_ and to create mutants in which the interaction is suppressed.

### Mutations in the GGDEF-FimV interaction surface alter PdeB localization dynamics

To further determine whether the decreased affinity between several GGDEF_PdeB_ variants and FimV_HubP_ affects polar recruitment of PdeB by HubP, we performed *in vivo* localization studies by fluorescence microscopy. To this end, the appropriate mutations were introduced into the chromosome of a *S. putrefaciens* strain producing functional fluorescently labeled PdeB-sfGFP (15). All substitutions in GGDEF_PdeB_ that negatively affected the interaction with FimV_HubP_ *in vitro* were stably produced at normal levels (**Figs. S4a-c**) but exhibited a significant decrease in polar localization *in vivo* (**Figs. 3a–d; Figs. S4d,e; Figs. S5a-f**). The PdeB variants bearing the G497A and K527E/Q528S substitutions within the degenerated I-site and the K578S substitution almost completely lost polar localization.

**Figure 3:**
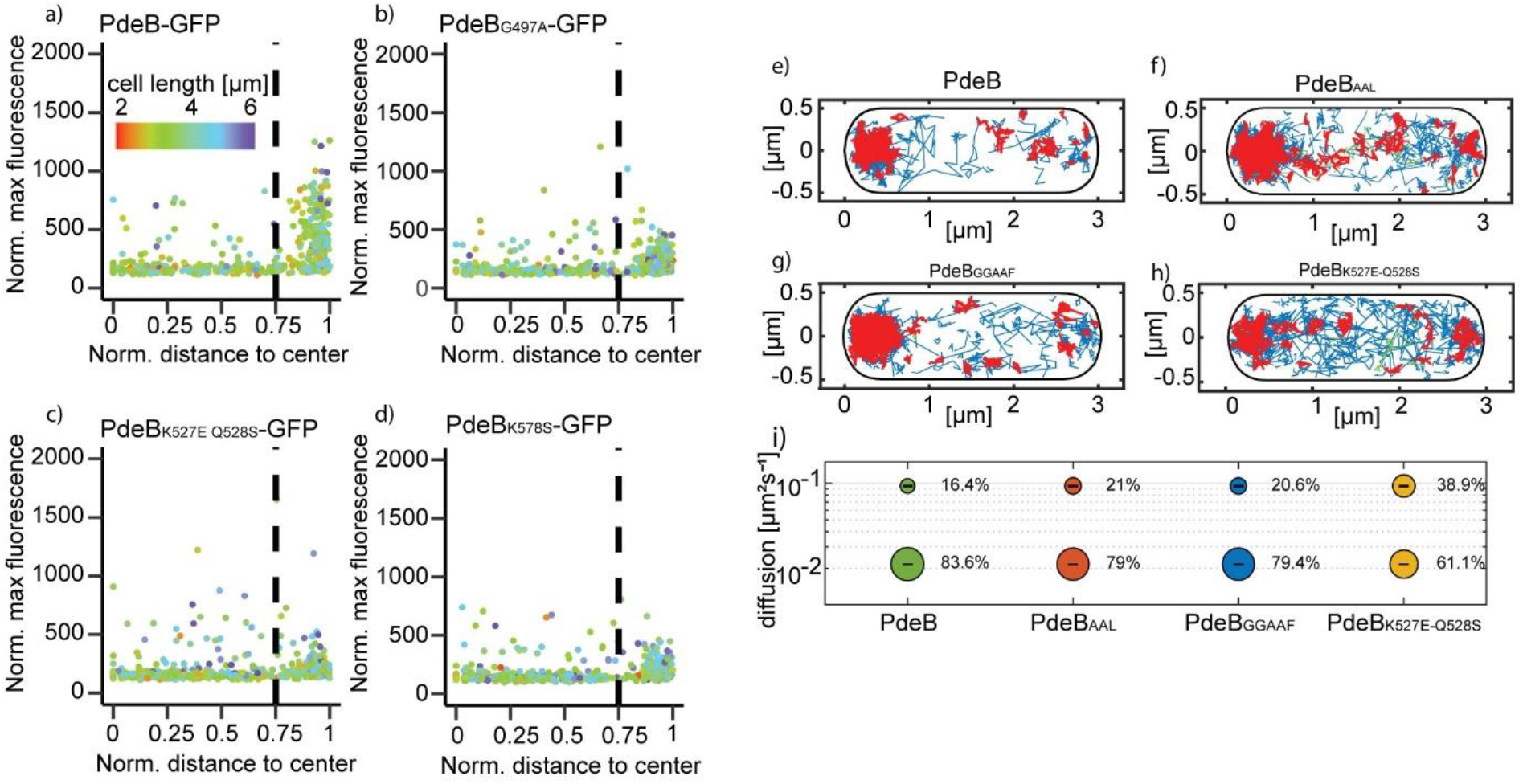
Localization behaviour of *Sp* PdeB-sfGFP variants. **a-d)** Residues that were shown to decrease the affinity of PdeB to HubP were genomically substituted in *S. putrefaciens pdeB-sfgfp*. The localization was quantified using BacStalk and is shown in 3d scatterplots where each dot represents an individual cell. Mutated proteins showed decreased polar localization compared to the wild-type version. **e – i)** The diffusion rates of mutants with mutations in the active center of the GGDEF and EAL domain, as well as in the interaction surface with HubP (h) were determined using single molecule microscopy of PdeB-mVenus. Projection of tracks into a standardized cell of 1 by 3 μm size. Blue trajectories represent freely diffusive molecules, red tracks molecules in confined motion (molecules staying within a radius of 120 nm, equal to three times the localization error), and green tracks represent molecules showing mixed behavior, i.e. switching between diffusive and confined motion. Bubble blot showing size and diffusion coefficients of determined populations. For a better comparison between PdeB and the mutant versions, diffusion constants were determined to best fit to all four proteins (0.095 μm^2^s^−1^ ± 0.001 for the slow population, 0.22 μm^2^s^−1^ ± 0 for the fast population).

To analyze the localization dynamics of PdeB in more detail, we monitored the intracellular diffusion of the protein *in vivo* using single molecule tracking (SMT) and determined the average time PdeB spent in static/slow moving (interacting with FimV_HubP_) or an unbound, freely diffusive mode. To this end, the C-terminal sfGFP tag to PdeB was replaced by mVenus. We observed that about 84% of the observed molecules were statically confined to the cell pole leaving only about 16% of the molecules that were freely diffusing (**Figs. 3e,i**). In a mutant bearing the K527E/Q528S substitution within GGDEF_PdeB_ the ratio shifted towards the freely diffusing subpopulation (61% static, 39% diffusing) (**Figs. 3h,i**). To further determine if the PdeB localization dynamic is affected by GTP binding (by the GGDEF domain) or the PDE activity (conferred by the EAL domain), we also determined the protein diffusion of PdeB-mVenus variants in which the GGDEF or the EAL domain were mutated by residue substitutions (GGAAF and AAL, respectively). The resultant PdeB-mVenus mutant versions exhibited only minor differences in diffusion compared to that of non-mutated PdeB-mVenus (**Figs. 3f,g,i**). Thus, PdeB is tightly bound to FimV_HubP_ via its GGDEF domain and therefore is almost exclusively present at the cell pole, and this localization pattern is unaffected by the enzymatic activity of PdeB.

### Substitutions within the GGDEF_PdeB_ interaction surface to FimV_HubP_ affect PdeB *in vivo* activity

The above results suggested that the interaction surface between GGDEF_FimV_ and FimV_HubP_ involves residues with a putative mechanistic function in the GGDEF domain, e.g., the salt-bridge forming lysine residue (K578) or the degenerated I-site (K527Q528; G497). We therefore hypothesized that interaction of the GGDEF domain of PdeB with FimV may result in differences in *in vitro* function of PdeB. To test this, we determined the intracellular c-di-GMP concentrations of *S. putrefaciens* wild-type cells and cells bearing substitutions in the residues in the GGDEF domain of PdeB that are critical for HubP/FimV interaction. We found that all substitutions resulted in significantly elevated c-di-GMP concentrations like that of a mutant completely lacking *pdeB* (**Fig. 4a**).

**Figure 4:**
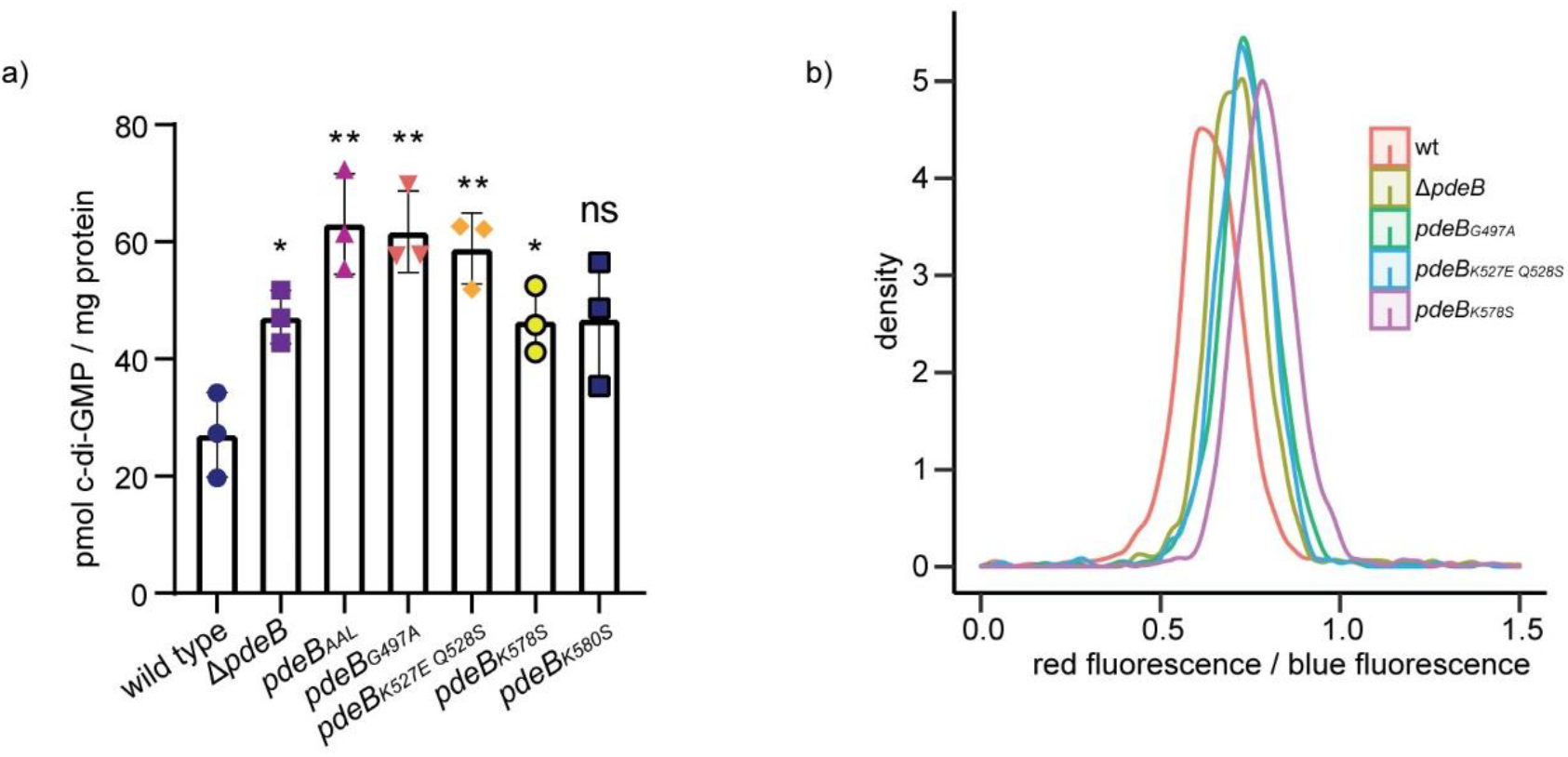
HubP affects the activity of PdeB *in vivo*. **a)** The cellular c-di-GMP of strains with reduced PdeB localization was extracted and quantified by mass spectrometry. Strains with reduced FimV/GGDEF interaction and polar localization of PdeB showed significantly increased c-di-GMP concentration. **b)** The c-di-GMP level of strains with reduced PdeB localization was determined using a fluorescence microscopy-based reporter assay. The tested mutants showed increased cellular c-di-GMP levels, indicating that polar localization of PdeB is required for full PDE activity.

To verify these findings at the single cell level, we adopted a plasmid-based c-di-GMP-responsive fluorescent reporter system for *Shewanella*, which relies on three c-di-GMP-dependent riboswitches mediating the production of destabilized turboRFP_AAL_ (32, 33). Additionally, a second fluorophore is constitutively produced that allows normalization of the c-di-GMP-derived signal to the plasmid copy number (**Figs. S6a,b**). In accordance to our *in vitro* results, all cells lacking *pdeB* or producing PdeB mutated in the GGDEF/FimV interaction surface displayed a higher concentration of c-di-GMP than wild-type cells (**Fig. 4b**). The results demonstrate that the residues within the GGDEF/FimV interaction surface are critical for *in vivo* function of PdeB and strongly suggest that interaction of PdeB with HubP at the cell pole stimulates PDE activity.

### PdeB promotes heterogeneity in c-di-GMP concentrations

As shown above, the c-di-GMP-responsive reporter system revealed that the *S. putrefaciens* wild type exhibits a, on average, lower level of c-di-GMP than mutants lacking or being mutated in PdeB. Of note, we also observed a broader distribution of c-di-GMP levels within the population, including a subpopulation with a very low c-di-GMP content (**Fig. 5a**). Thus, the presence and activity of PdeB leads to an increased heterogeneity of c-di-GMP levels of the cells within a population. We reasoned that part of the effect likely stems from the bimodal distribution of FimV_HubP_ during cell division (15). In addition, during single molecule tracking we observed a very low number of PdeB-mVenus tracks, and many cells completely lacked fluorescent signals. To determine the number of PdeB-mVenus molecules per cell, we quantified the photon count of single bleaching steps (i.e., single-fluorescent-protein bleaching) toward the end of the signal acquisitions. The total fluorescence intensity at the beginning of the acquisition (not yet bleaching) was then divided by that of the single fluorophores, relative to the cell size (6). By this, we found that about 25% of the cells did not contain any PdeB molecule, while the residual cells harbored one to forty PdeB molecules, most cells possessed one to seven copies (**Fig. 5b; Fig. S7**). Thus, in addition to the spatial distribution of active PdeB molecules per cell occurring during cell division further heterogeneity in c-di-GMP levels is mediated by significant differences in PdeB molecule numbers.

**Figure 5:**
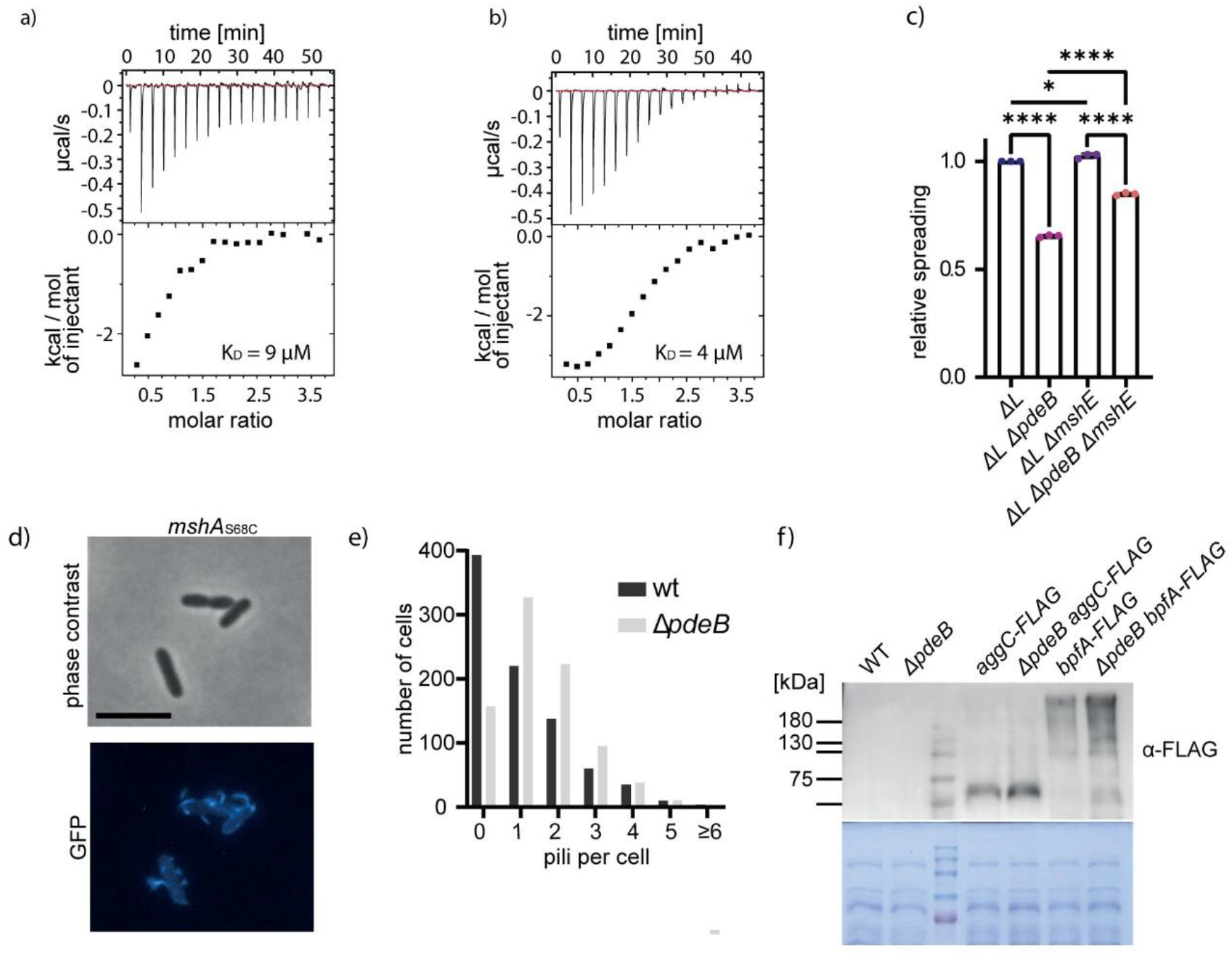
PdeB regulates *S. putrefaciens* adhesion factors. **a-b)** Binding of c-di-GMP to MshE (a) and its N-domain (b) was shown with ITC experiments. Both versions bind the nucleotide messenger with high affinity with a dissociation constant in the low micro molar range. **c)** The influence of MshE on the spreading phenotype of *ΔpdeB* was tested using soft agar motility assays. Since we previously showed that PdeB regulates the lateral flagellar system (15), these experiments were done in strains lacking the genes for the lateral flagellins. **d-e)** The number of extended MSHA pili in presence and absence of *pdeB* were quantified using fluorescence microscopy. The scale bar represents 5 μm. Significantly more extended MSHA pili were observed in the absence of *pdeB*, as shown in the histogram (e). **f)** Absence of *pdeB* also leads to increased amounts of Bpf proteins, as shown by immunoblot assays. The amount of two different proteins, BpfA and BpfD, were measured in presence and absence of *pdeB*.

Previous studies on *P. aeruginosa* showed that an asymmetric division of surface-associated cells results in one daughter cells prone to stay attached due to high concentrations of c-di-GMP and one flagellated daughter cell with low c-di-GMP levels, which is more prone to detach and leave. This behavior was termed ‘Touch-Seed-and-Go’ (13). The strict polar localization of PdeB and its regulation of surface adhesion factors in *S. putrefaciens* strongly suggested that PdeB promotes a similar behavior. We therefore monitored and quantified the detachment behavior of dividing *S. putrefaciens* cells adhered to a surface. To distinguish between the daughter cell with or without PdeB, we used a strain in which PdeB was C-terminally labeled by sfGFP (**Fig. 5d**). We observed that in about 80% of the division events that one cell detaches after cell fission. When a PdeB-sfGFP cluster was visible in one cell (in about 50% of the events), it was always this cell detaching (**Fig. 5c**). In a strain in which the PDE activity of PdeB-sfGFP was disrupted by introduction of a mutation within the EAL active center (PdeB-sfGFP_AAL_) a much smaller fraction of cells detached after finishing cell division (about 20%). Among these detaching cells, only about a 30% had a visible PdeB-sfGFP_AAL_ cluster.

Taken together, our data shows that, in *S. putrefaciens*, abundance and localization of PdeB promote heterogeneity in cellular c-di-GMP concentration within the population including an asymmetry between dividing cells, facilitate ‘Touch-Seed-and-Go’ behavior and thus regulate the spreading and proliferation of the population.

### PdeB regulates the abundance and activity of major surface adhesion factors in *S. putrefaciens*

Our results show that the activity of PdeB governs *S. putrefaciens* surface adhesion. However, so far it is unknown which surface adhesion factors are directly or indirectly regulated by PdeB. We therefore determined, whether major surface adhesion factors of *S. putrefaciens*, the surface adhesion protein BpfA (34–36) and the mannose-sensitive hemagglutinin (MSHA) type IV pilus (37, 38), are regulated by PdeB.

In the *Vibrio cholerae* MSHA pilus system, filament extension and retraction can be modulated by direct binding of c-di-GMP to the pilus extension ATPase MshE, which affects near-surface swimming and motile to sessile transition (39–41). Amino acid sequence alignments indicated the presence of the conserved c-di-GMP binding motif at the N-terminal domain of *S. putrefaciens* MshE (**Fig. S8**). To determine potential c-di-GMP binding of that region, full-length MshE as well as its predicted c-di-GMP-binding domain MshE-N (aa 2–145) from *S. putrefaciens* were purified (**Fig. S9**). Using isothermal titration calorimetry (ITC) and fluorescence polarization (FP) we found that both MshE and MshE-N bind c-di-GMP with high affinity (**Fig. 6a,b**). This data indicated that PdeB regulates MSHA pili extension via c-di-GMP concentrations. Therefore, we directly determined MSHA pili formation using a serine to cysteine substitution (S68C) within the major subunit of the pilus filaments MshA. The cysteine residue allowed covalent coupling of maleimide-ligated fluorophors to extended pili (41) and subsequent visualization and quantification of MSHA pilus formation by fluorescence microscopy (**Fig. 6d**). Compared to wild-type cells, mutants lacking *pdeB* displayed a significantly higher amount of extended MSHA pili (**Fig. 6e**). In addition, cells lacking MSHA pili showed increased spreading through soft agar, partially complementing a *ΔpdeB* mutant in polarly flagellated *S. putrefaciens* cells (**Fig. 6c**).

**Figure 6:**
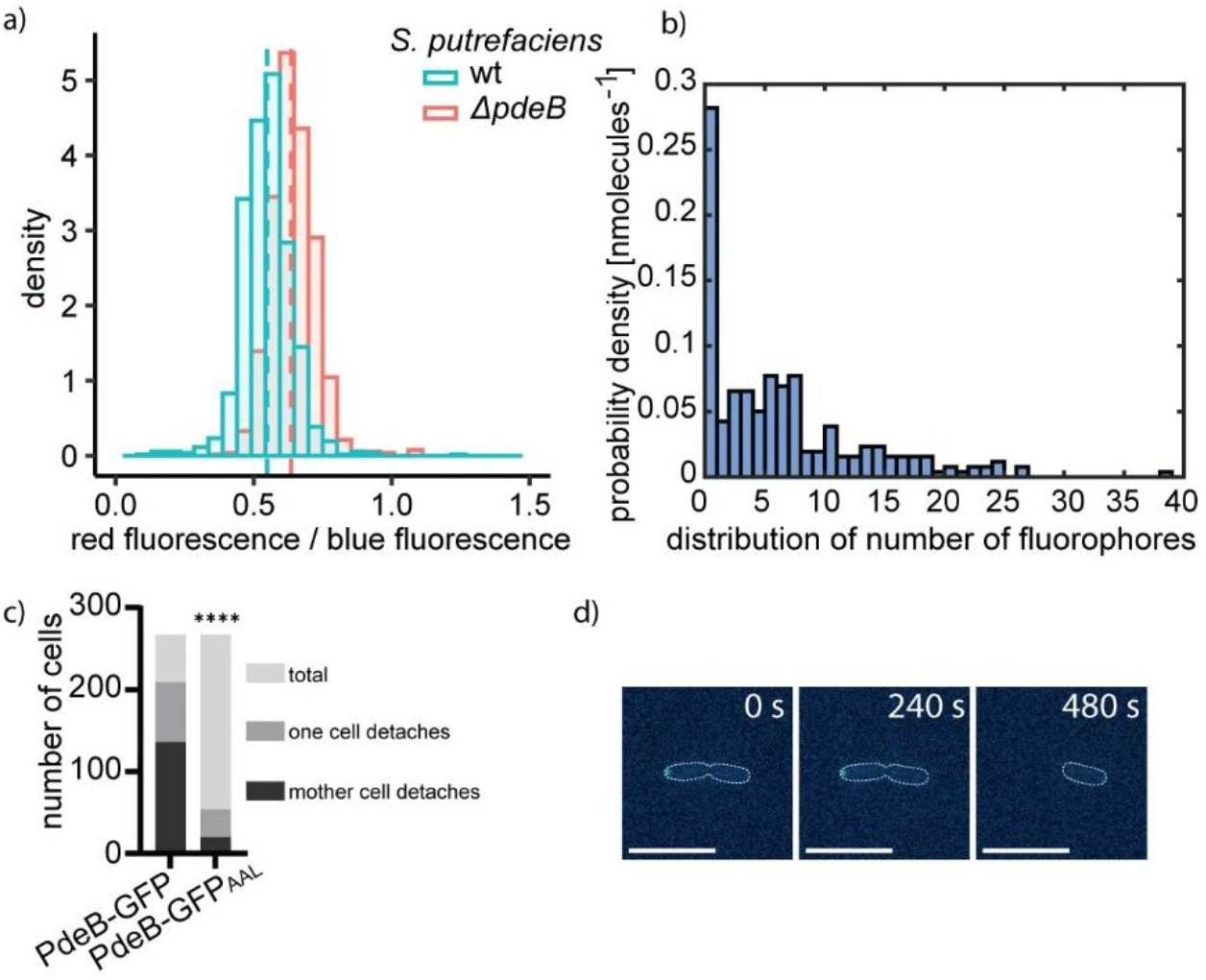
Polar localization of PdeB leads to heterogeneity. **a)** The c-di-GMP level of single cells in presence and absence of *pdeB* was measured using a fluorescence microscopy-based reporter assay. The obtained data was normalized to the plasmid number and the influence of *pdeB* on the heterogeneity was tested in *S. putrefaciens* (see **Fig. S6**). Absence of *pdeB* resulted in increased c-di-GMP levels. The histograms show that in presence of *pdeB* an additional subpopulation with low c-di-GMP levels is observed, likely due to asymmetric cell division. **b)** The number of PdeB-mVenus molecules in *S. putrefaciens* was measured using single molecule microscopy and is shown as histogram. Around 25% of the observed cells do not harbor PdeB-mVenus molecules. **c,d)** Polar localization leads to phenotypic heterogeneity and leads to the formation of a motily mother cell and a sessile daughter cell. 250 cell divisions were observed for each strain and results are shown in stacked bar plots (c). In presence of *pdeB* a motile mother cell and a sessile daughter cell is created. An exemplary time lapse is shown in (d), scale bar = 10 μm.

To investigate a potential effect of PdeB on the *S. putrefaciens* Bpf surface adhesion protein system, we determined the abundance of proteins BpfA and AggC (Sputcn32_3592) in the presence or absence of PdeB. To this end, the corresponding genes on the chromosome were mutated to encode protein versions with a C-terminal FLAG-tag, which allowed the quantification of the protein levels by Western blotting. All tested proteins showed a significantly higher abundance in the absence of PdeB (**Fig. 6f**). The results demonstrate that PdeB regulates the presence and activity of major c-di-GMP-dependent surface adhesion factors, i.e. the MshA pilus and the Bpf surface protein system, and by this governs cell-surface interactions which in turn affects flagella-mediated swimming and spreading through structured environments.

## Discussion

Previous studies have provided evidence that the polar activity of enzymes involved in c-di-GMP synthesis and turnover affects important cellular processes, such as cell cycle regulation in *C. crescentus* (9) or mediating the dynamics of a surface-associated or planktonic life style in *P. aeruginosa* (11–13). In these systems, bimodal distribution of c-di-GMP during cell division is usually controlled by abundance (e.g. through spatiotemporally regulated degradation) or specific activation of the corresponding proteins (e.g. through phosphorylation) (1, 12). Here we find that the *S. putrefaciens* GGDEF_PdeB_ domain serves as a spatial on-switch regulator for PdeB upon direct interaction with the FimV domain of the polar marker protein HubP, thereby restricting full PdeB activity to the flagellated cell pole.

### PdeB function depends on interaction with FimV

As in PdeB, in about two thirds of the proteins containing an EAL domain, this domain occurs as a GGDEF-EAL pair at the C-terminal end of the corresponding protein (42). In contrast to some of these hybrid proteins (43–45), PdeB does not function as a DGC *in vivo* or *in vitro* under the conditions tested. This is at least partly due to a glutamate residue conserved in PdeB orthologs (E467 in SpPdeB) replacing a non-variable functionally crucial aspartate in canonical GGDEF domains (30). We therefore conclude that the GGDEF domain of PdeB solely acts as a localizing and regulating domain.

Despite its inactivity as a DGC, GGDEF_PdeB_ is still able to bind GTP to its active center, which is crucial for PDE function. Previous work on different DGC-PDE hybrids (e.g., *Caulobacter crescentus* CC3396, *P. aeruginosa* RbdA and RmcA) demonstrated that GTP binding to an otherwise inactive GGDEF domain can regulate the downstream EAL domain’s PDE activity (46–49). Notably, the domain organization and structure prediction of PdeB closely resembles that of RbdA (**Fig. 2e**). Recent studies on RbdA indicate that GTP binding results in structural rearrangements of an RbdA dimer. These rearrangements release autoinhibitory interactions between the GGDEF and the EAL domains, which are then capable of binding and hydrolyzing c-di-GMP (48). Similarly, GTP binding to the GGDEF domains of an RmcA dimer is proposed to unlock and activate the EAL domains (49). The unlocking mechanism is based on a rearrangement of an alpha helix (hinge helix) that connects the DGC and PDE domains, which is also predicted to be present in PdeB. Based on the structural similarities, we hypothesize that PdeB phosphodiesterase activity is allosterically regulated by GTP in a similar fashion.

In addition to GTP binding, PdeB function *in vivo* requires the interaction with the C-terminal FimV domain of the polar landmark protein HubP. We found that tight interaction of the two proteins occurs between a negatively charged surface area of FimV and a positively charged surface area of the GGDEF_PdeB_ domain, which also includes the degenerated inhibitory (I-)site of the latter. Substitutions within these residues drastically lower the affinity between FimV and GGDEF_PdeB_ and shut down *in vivo* PDE function without affecting GTP binding. In functional dimers of canonical DGCs, c-di-GMP can bind to the I-site of one GGDEF domain and a second inhibitory site, the N-terminal R” site, of the second GGDEF domain. This locks the dimer in an inactive conformation in an end-product inhibition (30). However, as the R” site is absent in all PdeB orthologs, we hypothesize that FimV binding to GGDEF_PdeB_ does not immobilize the domains in the dimer but rather induces conformational changes that promote the formation of a complex competent for activation of the C-terminal EAL domains. Further structural studies are required to elucidate the exact underlying mechanism. Notably, the DGC DgcP Recently was shown to be directly recruited to the cell pole in *P. aeruginosa* by FimV, the orthologous polar landmark protein to HubP in *Pseudomonas* (50). DgcP affects surface adhesion/biofilm formation and twitching motility in *P. aeruginosa* and the DGC activity was decreased in cells lacking FimV. Thus, as PdeB, DgcP may also be activated through direct interaction with a polar marker protein, but so far direct evidence for this lacking. The GGDEF domains of DgcP and PdeB are not conserved in the critical residues required for PdeB recruitment and activation, which should be expected as both proteins possess opposite enzymatic activities.

In cells of *S. putrefaciens* and *S. oneidensis* PdeB mediates a considerably increased heterogeneity with respect to c-di-GMP concentrations in the presence of PdeB. This heterogeneity can be explained by at least two closely connected factors, (i) the strictly monopolar PDE activity in concert with a non-uniform appearance of PdeB at the new cell pole and (ii) the wide range of a comparably small number of PdeB copies per cell. Thus, by regulating flagella-mediated motility and at least two *Shewanella* c-di-GMP-responsive surface-adhesion factors PdeB promotes a wide range of cell behaviors with respect to spreading and attachment in each population. It is yet unknown how production of *Shewanella* PdeB is regulated to result in such a pronounced heterogeneity in abundance and timing of appearance after cell division.

### PdeB regulates the transition between sessile and planktonic lifestyle

As for many bacterial species, c-di-GMP-mediated signaling is crucial for regulating the switch from sessile to planktonic lifestyle in *Shewanella* sp., and PdeB has an important role in balancing planktonic and surface-associated states of the cells. In *S. putrefaciens*, PdeB stimulates the formation and activity of the lateral flagella system, which positively affects swimming and spreading of the cells (15, 51, 52). The effect on the lateral flagella is likely mediated by the c-di-GMP-responsive transcriptional flagellar regulator FlrA2 and the c-di-GMP-binding flagellar motor regulator MotL (15, 53, 54). Here we demonstrated that PdeB also negatively regulates abundance and activity of two major factors that directly mediate cell-surface interactions, the surface adhesion protein BpfA and the mannose-sensitive hemagglutinin (MSHA) type IV pilus. The MSHA pilus system has previously been demonstrated to mediate cell-surface interactions of *S. oneidensis* (37, 38) and to affect the ability of this species to colonize the Zebrafish gut (55). Here we found that the MSHA pili negatively affect the spreading of *S. putrefaciens* through soft agar, which accounts for a large part of the PdeB-related spreading phenotype of the polar flagella of *S. putrefaciens* and likely also in *S. oneidensis*. In *Vibrio cholerae*, the pilus extension ATPase, MshE, is stimulated by binding c-di-GMP (40, 56), and active *Vc*MSHA pili also limit cell spreading through polysaccharides and assist in initiating surface adhesion by actively swimming cells (39, 41). Our results strongly suggest that the *S. putrefaciens* MSHA pilus system functions in a similar fashion. The BpfA surface protein and the corresponding transport and control system of *Shewanella* are largely homologous to the well-characterized Lap system of *Pseudomonas fluorescens* (reviewed in (57)) and have an important role in mediating cell-surface interactions (34, 36). In *S. putrefaciens*, BpfA production and cell-surface abundance are regulated by c-di-GMP concentrations, e.g., via the flagellar master regulator FlrA_1_ and the oxygen-responsive DGC DosD (35, 53); our results demonstrate additional regulation of BpfA abundance by PdeB. Notably, both LapA and well as *Vc*MSHA pili are thought to be involved in mediating c-di-GMP-dependent attachment and detachment of cells to a substratum (41, 57), which we now propose to similarly occur for the corresponding *Shewanella* systems.

The signal PdeB is responding to is yet unknown. In *P. aeruginosa* the PDE Pch is recruited by the main chemotaxis histidine kinase CheA, which in addition governs Pch activity by phosphorylation (12). By this, *P. aeruginosa* elegantly links the perception of a wide range of external signals via the chemotaxis sensor array to cellular behavior in planktonic as well as in surface-associated cells. The asymmetry of c-di-GMP levels in dividing cells mediated by Pch also affects the surface attachment/detachment response of *P. aeruginosa*, which depends on polar recruitment of the c-di-GMP-responsive pilus regulator FimW upon surface-association (13). In *S. putrefaciens* and *S. oneidensis*, PdeB appears to be constantly but heterologously produced irrespective of the media conditions (15), and polar recruitment of PdeB occurs independently of surface attachment. However, PdeB appears to be solely active in complex media (14, 15). Our data strongly suggests that PdeB activation requires GTP binding and may therefore be dependent on the cellular GTP concentrations. As previously discussed, the cellular GTP level is correlated with that of the alarmone (p)ppGpp and thus also reflects the cellular nutrient levels, so that PdeB activity is shut down under conditions of low nutrients (46, 48). In addition, PdeB possesses two putative sensor domains, the N-terminal periplasmic domain and the cytoplasmic PAS domain. It has been suggested that sensor PDE proteins (e.g. RmcA and RbdA) may function as rheostats that integrate several signals to tune the proteins activity (48, 49) similar to the sum of different signals transmitted by the chemotaxis array to Pch of *P. aeruginosa* via CheA. The periplasmic domain of PdeB has no striking homology to other sensing domains but is required for full PDE function (15). The domain’s localization hints at an external signal, such as a small molecule or peptide. Notably, compared to the remaining protein, this hypothetical sensing domain is less well conserved among *Shewanella* sp. As members of this genus occur in wide range of different environments (58), it is conceivable that *Shewanella* PdeB orthologs have evolved to respond to different environmental signals. The second potential sensing domain of PdeB, the cytoplasmic PAS domain, is also required for full PdeB function (15), and PAS domains could be involved in the perception of a wide range of potential signals (59). As PdeB orchestrates population heterogeneity and behavior, ongoing current studies are directed at identifying the yet elusive signal(s).

## Materials & Methods

### Strains, growth conditions and media

The strains used are listed in **Table S1**. *Shewanella* strains were routinely cultured either at room temperature or 30°C in LB (10 g l^−1^ tryptone, 5 g l^−1^ yeast extract, 10 g l^−1^ NaCl, pH 7) or lactate (LM) medium (10 mM HEPES, pH 7.5, 200 mM NaCl, 0.02% (w/v) yeast extract, 0.01% (w/v) peptone, 15 mM lactate). *Escherichia coli* strains were standardly grown in LB medium at 37°C, if not indicated otherwise. When appropriate, media were supplemented with 50 μg·ml^−1^ kanamycin or 10% (w/v) sucrose. Cultures of the *E. coli* conjugation strain WM3064 were supplemented with 2,6-diaminopimelic acid (DAP) to a final concentration of 300 μM. For plates, the appropriate medium was supplemented with 1.5% (w/v) agar.

### Plasmid and strain constructions

Plasmids are listed in **Table S2** and were generated essentially as described in (15). The used oligonucleotides are listed in **Table S3**. Plasmids were constructed by PCR amplification of the DNA fragments of interest and subsequent Gibson assembly with the appropriate vector backbone (60). All plasmids were verified by sequencing. To modify the genome of *Shewanella* species flanking regions of 500 to 750 nucleotides upstream and downstream of the integration site were amplified by PCR and inserted into the suicide vector pNTPS‐138‐R6K (61). The resulting plasmids were transferred into *Shewanella* from *E. coli* WM3064 by conjugation. Correctly mutated strains after sequential homologous recombination were identified by plating the cells on either kanamycin- or sucrose-containing LB medium plates. Successful integration or deletion was verified using colony PCR.

### Soft agar motility assay

Overnight LB cultures of the appropriate strains were diluted to an OD_600_ of 0.02 and grown to mid exponential growth phase (OD_600_ of 0.5). 2 μl of respective culture were dropped on LB soft-agar plates (0.25% (w/v) agar). The plates were incubated at room temperature overnight. The plates were scanned and the swim radius was quantified. Cultures of strains to be directly compared were always placed on the same plate.

### Fluorescence microscopy

The strains to be imaged were cultivated overnight in appropriate medium and then subcultivated until reaching exponential growth phase (OD_600_ of 0.2) or another desired OD_600_. Three microliters of culture were spotted on an agarose pad (LM medium solidified with 1% (w/v) agarose). Fluorescence images were recorded using a Leica DMI 6000 B inverse microscope (Leica, Wetzlar, Germany) equipped with an sCMOS camera and an HCX PL APO 100×/1.4-numerical-aperture objective using VisiView software (Visitron Systems, Puchheim, Germany). The obtained images were further processed using ImageJ (62) (https://imagej.nih.gov) to adjust contrast, to change grayscale to color, to define representative areas for display, and to add scale bars. Adobe Illustrator CS6 was used to add an outline of cells where required and to create the final assembly of panels. Cells from at least three independent cultures were imaged.

### Fluorescent labelling of MSHA-pili

To fluorescently label MSHA type IV pili, we adopted pili labeling protocols for *Vibrio cholerae* (41, 63) and flagella labeling in *S. putrefaciens* (64). Overnight cultures of the desired strains in LB were diluted to an OD_600_ of 0.02 and re-cultivated to the desired OD_600_ of 0.2. Cells were harvested by centrifuging 100 μl of each culture at 4000 rpm for 5 minutes at room temperature. The pellets were washed with 50 μl phosphate-buffered saline (PBS; NaCl, 137 mM; KCl 2.7 mM; Na_2_HPO_4_, 10 mM; KH_2_PO_4_, 1.8 mM; pH 7.4). 1.5 μl CF®488A Maleimide was added to reach final concentration of 25 μg · ml^−1^. The cells were incubated for 15 minutes in darkness to prevent bleaching. Afterwards the cells were again sedimented by centrifugation at 4000 rpm for 5 minutes at room temperature and washed with 100 μl PBS. The washing step was repeated and the cells were resuspended in 100 μl LM medium. 3 μl of the stained culture was dropped on a 1% (w/v) LM-agar pad for microscopy.

### Quantification of polar PdeB-sfGFP variants by fluorescence microscopy

To quantify the signal intensity of polarly localized PdeB-sfGFP variants, overnight cultures were subcultivated as described above and grown at 30 °C until OD_600_ of 0.5 was reached. Aliquots were then used for microscopy as described. The obtained images were analyzed using the BacStalk software (Hartmann et al, 2020) to obtain data for the cell length, mean fluorescence, maximum fluorescence and distance of the maximum fluorescence to the cell center. Following, at least 300 cells were selected at random and a new parameter was calculated called “normalized distance to center” by using following formula:

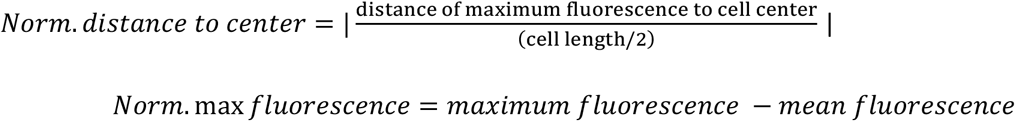

The cluster intensity was normalized by subtracting the mean fluorescence from the maximum fluorescence. Both values were then plotted against each other where each point represents an individual cell. The color of each point represents the cell length of respective cell.

### c-di-GMP *in vivo* fluorescence reporter assay

The c-di-GMP reporter plasmid pMMB-Gm-Bc3-5 AAV was kindly provided by the group of Fitnat Yildiz (33). The plasmid was introduced in *Shewanella* via conjugation. Overnight cultures supplemented with 5 μg · ml^−1^ gentamycin were diluted to an OD_600_ of 0.02. Cultures were grown to approximately OD_600_ 0.6, diluted and 3μl were dropped on LM-agar pads. Pictures with exposure time of 100 ms were taken using the microscopic set-up described above. The relative fluorescence intensity (RFI) was then determined by obtaining the ratio of turboRFP/AmCyan as quantification of the amount of intracellular c-di-GMP. The fluorescence images were analyzed using BacStalk (65) and parameters such as cell length, maximum fluorescence intensity, mean fluorescence intensity and distance of maximum fluorescence to cell center were obtained. For downstream analysis we selected at least 500 cells by random with a cell length between 2 μm and 6 μm.The obtained data was then plotted using the programming environment R and the software Rstudio (Rstudio Inc.). To verify that the reporter system is suitable for the used *Shewanella* species, we plotted the red fluorescence intensity against the blue fluorescence intensity and determined the expected linear correlation (**Fig. S6**). The quotient of red fluorescence divided by blue fluorescence was used as parameter for the intracellular c-di-GMP level and was plotted against the cell length as additional control (**Fig. S6**).

### Attachment/detachment assay of *S. putrefaciens* cells

Overnight cultures of *S. putrefaciens pdeB-sfGFP* and *pdeB-sfGFP_AAL_* in LM were diluted with fresh LM to an OD_600_ of 0.02, and the cultures were further grown at room temperature for 3-4h. After diluting the culture 1:200, an appropriate aliquot was filled into microscopy dishes (ibidi, München, Germany, cat.no 81156), that allow microscopic observation of cells attached to the bottom of the dish. Time lapse images were taken every 40 s in the GFP and phase contrast channels at an exposure of 30 ms. The resultant image stacks were further processed using Fiji imageJ (62) (https://imagej.nih.gov). 250 dividing cells/cell pairs of each strain were counted and analysed.

### Single molecule tracking

Cultures were grown to an OD_600_ of 0.5-0.6 in LB medium. 5 μl of the culture were placed on a cover slide and cells were immobilized with an agar pad (1% (w/v) agarose in LB). The microscope used for the single molecule tracking was a Nikon Eclipse Ti microscope (100x oil-immersion objective, NA=1.49). Cells were imaged with the central part of a 514 nm-laser (TOPTICA Beam Smart, maximum power = 100 mW), using about 160 to 200 W cm^−2^; single molecule level was reached rapidly after 100 frames. Movies with 2000 frames were taken with an EM-CCD camera (ImageEM X2 EM-CCD, Hamamatsu). Movies were cropped so that the first 100 frames were removed, performed in Fiji software (66). Cell outlines were determined by the software Oufti (67), and afterwards trajectories were determined with the software UTrack 2.2.1 (68). SMTracker 1.5 (6) was used for all further the analyses of protein dynamics. Diffusion constants were determined using squared displacement (SQD)-analyses. Trajectories were categorized into confined motion, for which a radius of 120 nm was set, or as freely diffusive, or having mixed behavior.

### Fluorescence-based quantification of molecule number

For molecule quantification, movies of PdeB-mVenus-producing cells, or of cells lacking any mVenus fusion (for determination of auto-fluorescence), were acquired in the same medium, on the same day. 11 complete movies (including the first 100 frames) of each strain were taken and for each strain; one video of the empty slide was captured for background correction. The analysis was carried out as described in SMTracker 2.0 (see https://sourceforge.net/projects/singlemoleculetracker/) was used for the determination. Initial fluorescence in cells was divided by determined single bleaching steps of PdeB-Venus, to calculate numbers of fluorophores per cell (for which the length is known from Oufti software, see above) and thereby average molecule number.

### Protein expression and purification

The plasmid pET24c was used for heterologous protein expression in *E. coli* BL21(DE3) (New England Biolabs, Frankfurt, Germany). Cells were grown in LB medium supplemented with 50 μg ml^−1^ kanamacin to an OD_600_ of 0.75 at 37°C and vigorous shaking. Prior to induction with d-(+)-lactose-monohydrate (12.5 g liter^−1^) cells were chilled for 10 minutes in an ice bath and expression was carried out overnight at 16°C under vigorous shaking. The cells were harvested the next day by centrifugation (5,000 rpm, 10 minutes, 4°C) and the resulting cell pellet was stored at −20°C until further use. Frozen cell pellets were resuspended in the appropriate buffer required for downstream applications supplemented with 0.2 % Tween20 and 40 mM imidazole and lysed by sonification (Bandelin Sonoplus). The resulting lysate was centrifuged for 30 minutes at 20,000 rpm to remove cell debris and the supernatant was filtered with a 0.45 μm filter before loading onto an equilibrated 5 ml His Trap HP column (GE healthcare) connected to an ÄKTA pure 25 system. Unspecifically bound proteins were removed by washing with 10 column volumes (CV) lysis buffer and proteins were eluted with a linear gradient of buffer supplemented with 600 mM imidazole. Eluted proteins were further purified and analyzed by size exclusion chromatography (SEC). The HiLoad Superdex 200 pg column (GE Healthcare, Chicago) was used for preparative runs while the Supedex200 increase column (GE Healthcare, Chicago) was used for analytical and semipreparative runs. Respective SEC columns were equilibrated with the appropriate assay buffer required for downstream applications without other additives. Proteins were isocratically eluted and the concentration was determined using a spectrophotometer (NanoDrop Lite, Thermo Scientific) before snap freezing in liquid nitrogen and storage at −80 °C. Protein size was analyzed using a calibration curve of proteins included in the high molecular weight calibration kit (GE Healthcare, Chicago). To test frozen samples for aggregation, SEC was used as described above. To purify protein complexes, both interaction partners were purified using affinity chromatography. Fractions containing the two proteins of interest were then mixed and incubated for at least 1 hour at 4 °C. The sample was then further purified using SEC as described above to remove unbound proteins.

### Protein pull down assays

The interaction of GGDEF_PdeB_ variants with FimV_HubP_ were tested using amylose resin based pull down assays. The purified MBP-tagged GGDEF domains (10 μM) were mixed equimolarly with purified FimV and then incubated overnight with equilibrated amylose resins (New England biolabs, UK). The beads were then washed 5 times with high salt Tris-buffer (50 mM Tris-HCl, 500 mM NaCl, 100 mM KCl, 5 mM MgCl_2_, pH 8). Then, proteins were then eluted using SDS-PAGE sample buffer and the eluate was analyzed using SDS-polyacrylamide gelelectrophoresis (SDS-PAGE).

### Western Blotting

Protein fusions were tested for stability and expression using Western blot analysis. Cells were grown in liquid culture until exponential phase to generate lysates. The concentration was adjusted to OD_600_ of 10, and 10 μl were loaded onto an SDS-gel and immunoblot detection was performed as described previously (15). Polyclonal antibodies against GFP were used to detect proteins fused to sfGFP or mVenus. Respective antibodies harbored a HRP fusion, and SuperSignal West Pico chemiluminescent substrate (Thermo Scientific, Schwerte, Germany) was used to generate a luminescence signal. The signal was then detected using a Fusion‐SL chemiluminescence imager (Peqlab, Erlangen, Germany).

### Biolayer interference assays

To analyze the effect of in frame amino acid substitutions on the affinity of *So*FimV to *So*GGDEF, the dissociation constant K_D_ was determined by biolayer interference assays. *So*FimV was used as ligand and purified in HEPES buffer (20 mM HEPES, 250 mM NaCl, 50 mM KCl, 5 mM MgCl2) before biotinylation using EZ-Link NHS-PEG4-biotin (Thermo Fisher Scientific, Waltham, Massachusetts) in equimolar concentration. Free biotin was removed using zebra spin desalting spin columns (Thermo Fisher Scientific, Waltham, Massachusetts).

The assay was carried out using the BLITZ system (ForteBio, UK) with streptavidin biosensors. For experiments that determined only the association medium salt HEPES buffer (50 mM HEPES, 250 mM NaCl, 50 mM KCl, 5 mM MgCl2, 0.02% (v/v) Tween20, pH 7.5) was used. However, the ionic strength of medium salt buffer was not high enough to induce sufficient dissociation, and, therefore, high salt Tris-buffer (50 mM TrisHCl, 500 mM NaCl, 50 mM KCl, 5 mM MgCl_2_, 0.02% (v/v) Tween20, pH 8) was used for association-dissociation experiments. The sensors were hydrated for at least 10 minutes in assay buffer and runs started with 30 seconds equilibration until four μl of the ligand protein was bound to the sensor for two minutes followed by 30 seconds washing. Then the association of the analyte protein was measured in real time for 300 s. For association-dissociation kinetic experiments this step was followed by 300 s washing with high salt Tris buffer (50 mM Tris-HCl, 500 mM NaCl, 100 mM KCl, 5 mM MgCl_2_, 0.2% (v/v) Tween-20, pH 8) and the BLITZ PRO software was used to determine the kinetic constants. For each interaction pair this was repeated at least 8 times in increasing concentrations until the K_D_ value was stable. Data was further analyzed using the GraphPad PRISM software.

### MANT-GTP binding assays

To verify the functionality of mutated *So*GGDEF proteins they were tested for GTP binding using MANT-GTP (Jena Bioscience, Germany). Proteins were diluted to 1 μM and 100 μl of each sample was added in triplicates into a white round bottom 96-well plate before MANT-GTP with a final concentration of 1 μM was added. The plate was then incubated for 30 minutes at room temperature on a plate shaker before the fluorescence (excitation: 295 nm, emission: 448 nm) was measured in a Tecan Infinite M200 plate reader (Tecan, Switzerland).

### Fluorescence anisotropy assays

Fluorescence anisotropy was carried out using a FP-8300 fluorescence spectrometer (Jasco, Pfungstadt, Germany) with high precision cells (Hellma Analytics, Müllheim, Germany). The titration was performed in assay buffer and all samples were degassed with a MicroCal ThermoVac (Malvern Pananalytical, Kassel, Germany) immediately before the experiment. As a ligand, we used MANT-labelled c-di-GMP (Biolog Life Science, Bremen, Germany) or GTP (JenaBioscience, Jena, Germany) and kept the ligand concentration constant during the titration. For MANT-c-di-GMP, we titrated increasing volumes (0.5 μl, 0.5 μl, 1 μl, 2 μl, 4 μl, 8 μl, 16 μl, 32 μl, 64 μl, 128 μl) of a solution consisting of 100 μM protein and 0.7 μM MANT-c-di-GMP to 60 μl 0.7 μM MANT-c-di-GMP at 25 °C. For MANT-GTP, we titrated increasing volumes (0.5 μl, 0.5 μl, 1 μl, 2 μl, 4 μl, 8 μl, 16 μl, 32 μl, 64 μl, 128 μl) of a solution consisting of 100 μM protein and 1 μM MANT-GTP to 60 μL 1 μM MANT-GTP at 25 °C. After each titration step, the sample was properly mixed, the change in fluorescence anisotropy measured five times (λ_exc_ 355 nm, λ_em_ 448 nm) and the results averaged. The data points are the mean of at least two replicates ± standard deviation. The data was fitted using the following quadratic binding equation for a one-site specific binding model (Graph Pad Prism 5). The K_D_ are given as fit ± standard deviation.

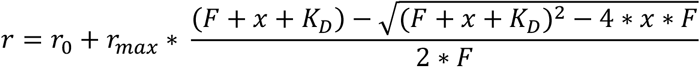

R = anisotropy, r_0_ = anisotropy before titration, r_max_ =maximum anisotropy, F = fluorophor concentration, x = protein concentration, K_D_ = dissociation constant

### Isothermal titration calorimetry assays

Isothermal titration calorimetry (ITC) was performed with MicroCal ITC2000 (Malvern Pananalytical, Kassel, Germany). The proteins were rebuffered five times into a tenfold volume of assay buffer and then concentrated using Vivaspin 6, 10,000 MWCO (Sartorius, Göttingen, Germany) at 4900 × g and 4 °C. The rebuffered samples were degassed immediately before ITC with MicroCal ThermoVac (Malvern Pananalytical, Kassel, Germany). The experimental setup was as follows: 18 injections of 2.0 μl 50 μM c-di-GMP were added to 200 μl 50 μM protein solution with an injection time of 4 s and 180 s spacing between each injection at 25 °C, and with constant stirring at 750 rpm. The first injection was set to 0.4 μl to remove air and mixed reactants from the tip. For each experiment, we performed a ligand-to-buffer (LtB) titration as well as a buffer-to-protein (BtP) titration with the same experimental setup to correct for possible heat of dilution introduced by the ligand or protein. The data was analysed using MicroCal Analysis (OriginLab). The first injection peak was discarded and the isotherms of the LtB and BtP controls were subtracted from the experimental isotherm. We then performed a nonlinear fit with one set of binding sites.

### iSCAM assays

Proteins were purified in low salt HEPES buffer (20 mM HEPES, 50 mM NaCl, 50 mM KCl, 5 mM MgCl2, pH 7.5) as described above using Ni-NTA affinity chromatography and SEC. Mass spectra were acquired on a Refeyn OneMP mass photomoeter (Refeyn, UK). The instrument was calibrated with Native Mark standards (ThermoFisher) at room temperature before use. We used high precision glass coverslips, 24×50 mm^2^, No. 1.5H, (Marienfeld, Germany) together with self-adhesive CultureWellTM gaskets (Grace Bio-Labs). Before measurements, samples were prepared at 1 μM. Before each measurement, the instrument was focused using 18 μl of room temperature buffer. 2 μl of were then added to the drop and mixed before measurement. We acquired 6000 frames for each measurement. Movies were analyzed using the DiscoverMP software (Refeyn, UK).

### Hydrogen/deuterium exchange mass spectrometry (HDX-MS)

HDX-MS experiments were conducted similarly as described (69). In brief, 5 μl of 60 μM concentrated individual FimV or the FimV/MBP-GGDEF_PdeB_ complex were mixed with 45 μl of D_2_O-containing buffer (20 mM HEPES-Na, pH 7.5, 200 mM NaCl, 20 mM KCl, 20 mM MgCl2) and incubated at 25 °C. After 0.25, 0.5, 1, 2 or 10 min, 50 μl of ice-cold quench buffer (400 mM KH_2_PO_4_/H_3_PO_4_, pH 2.2) were added and the mixture injected into an ACQUITY UPLC M-Class System with HDX Technology (Waters) (70). Undeuterated samples were obtained similarly by mixing with H2O-containing buffer. Samples were digested online with a column containing porcine pepsin (Enzymate Pepsin Column, 300 Å, 5 μm, 2.1 mm x 30 mm (Waters)) at a flow rate of 100 μl min^−1^ in H_2_O + 0.1% (v/v) formic acid at 12 °C, and the resulting peptides trapped for 3 min on an AQUITY UPLC BEH C18 1.7 μm 2.1 x 5 mm VanGuard Pre-column (Waters) kept at 0.5 °C. Subsequently, the trap column was placed in line with an ACQUITY UPLC BEH C18 1.7 μm 1.0 x 100 mm column (Waters) and the peptides separated at 0.5 °C with a gradient of H2O + 0.1% (v/v) formic acid (eluent A) and acetonitrile + 0.1% (v/v) formic acid (eluent B) at 30 μl min^−1^ flow rate as follows: 0-7 min/95-65% A, 7-8 min/65-15% A, 8-10 min/15% A. Peptides were ionized by electrospray ionization (capillary temperature 250 °C, spray voltage 3.0 kV) and mass spectra acquired over 50 to 2,000 m/z on a Synapt G2-Si HDMS mass spectrometer with ion mobility separation (Waters) in HDMS^E^ or HDMS mode for undeuterated and deuterated samples, respectively (71, 72). Lock mass correction was conducted with [Glu1]-Fibrinopeptide B standard (Waters). During separation of the peptides, the pepsin column was washed three times with 80 μl of 4% (v/v) acetonitrile and 0.5 M guanidine hydrochloride and blanks were performed between each sample. Three technical replicates (independent HDX reactions) were measured per incubation time point. Peptide ions were identified with ProteinLynx Global SERVER (PLGS, Waters) from the non-deuterated samples as described previously (73) and matched to peptides with a database containing the amino acid sequences of FimV, porcine pepsin and their reversed sequences. Deuterium incorporation into peptides was quantified with DynamX 3.0 (Waters) as described (73).

### Protein Cross-linking

The used proteins were purified by NiNTA affinity chromatography in HEPES buffer (20 mM HEPES, 250 mM NaCl, 50 mM KCl, 5 mM MgCl_2_). Fractions containing the proteins of interest were then separated into two batches before SEC were one sample was further purified in HEPES buffer and the other in PBS. The next day, 10μM proteins were crosslinked by addition of 1mM DSBU and samples were incubated for 30 minutes at room temperature before the crosslinking reaction was stopped by adding Tris-buffer to a final concentration of 10 mM. The crosslinking was verified by SDS-PAGE. Crosslinked proteins were digested by the addition of 1μg trypsin (Promega) and overnight incubation at 30°C. A final of 0.5 % sodiumdeoxycholate was spiked to the samples followed by reduction (Tris-2-Carboxyethyl phosphine, 5mM, 90°C, 10 min) and alkylation (Iodoacetamide, 10mM, 25°C, 30min), and an additional digestion step using 1μg trypsin for 2 h. Finally, peptides were desalted using C18 reversed phase solid phase extraction cartridges (Macherey-Nagel).

### LC-MS/MS analyses and identification of crosslinked peptides

LC-MS/MS analysis of digested crosslinked protein complexes was performed on Q-Exactive Plus mass spectrometer connected to an electrospray ionization source (Thermo Fisher Scientific). Peptide separation was carried out using a Ultimate 3000 nanoLC-system (Thermo Fisher Scientific), equipped with a C18 resin column (Magic C18 AQ 2.4 μm, Dr. Maisch) packed in-house. The peptides were first loaded onto a C18 precolumn (preconcentration set-up) and then eluted in backflush mode with a gradient from 98% solvent A (0.15% (v/v) formic acid) and 2% solvent B (99.85% acetonitrile, 0.15% formic acid) to 35% solvent B over 60 min. The flow rate was set to 300 nl min^−1^. The data acquisition mode for the initial LFQ study was set to obtain one high-resolution MS scan at a resolution of 70,000 (*m/z* 200, MS1) and 17,500 (*m/z* 200, MS2) with scanning range from 375 to 1,500 *m/z* followed by MS/MS scans of the 10 most intense ions. To increase the identification efficiency of MS/MS attempts, the charged state screening modus was adjusted to exclude unassigned, singly, and doubly charged ions. The dynamic exclusion duration was set to 30 s. The ion accumulation time was set to 100 ms (both MS and MS/MS). The automatic gain control (AGC) was set to 3 × 10^6^ for MS survey scans and 1 × 10^5^ for MS/MS scans. Collision was induced by high collision dissociation (HCD) with an NCE 23. MS raw files were converted into mgf files and analyzed using MeroX (v2.0) (74) in default settings.

### *In silico* protein analyses

*Protein sequence alignments and position based weight maps.* The NCBI and Pfam database were accessed to obtain the amino acid sequences of PdeB homologues and other GGDEF domains. Alignments were created using the Clustal omega tool (75) and were further analyzed with JalView (76). The position based weight map of PdeB homologues from different *Shewanella* species was created with the Seq2Logo tool (77) using the P-Weighted Kullback-Leiber clustering algorithm.

*Protein structure prediction and visualization.* The protein structure of PdeB (PAS-GGDEF-EAL) was predicted using the Phyre2 tool (78), while the structure of the C-terminal region of HubP was modeled using the Swiss Model algorithm (Waterhouse et al, 2018). The obtained data was then analyzed using Pymol (The PyMOL Molecular Graphics System, Version 2.0 Schrödinger, LLC) and the included APBS electrostatics tool.

### Extraction and analysis of cyclic dinucleotides

Extraction and quantification of nucleotide messengers was conducted as described in (79). Briefly, cells were grown to exponential phase before harvesting by centrifugation. PDEs and DGCs were inactivated by heating for 10 minutes at 95 °C and nucleotides were then extracted using a mixture of acetonitrile, methanol, and water (2:1:1). The protein amount was quantified using BCA assays with the residual pellet and extracted nucleotides were dried using vacuum centrifugation and stored at −20 °C before further use. To quantify the extracted nucleotides, mass spectrometry was operated in positive ionization mode. MS was performed with 50 μl each of calibrators, quality controls and biological samples and analyte separation was done accordingly to the HPLC gradient method. The obtained data of the calibrators was used to generate a calibration curve and the MS/MS signals obtained by the biological samples were analyzed by calculating the peak areas and calculating the ratios compared to the calibration signals to obtain precise measurements of the nucleotide concentrations.

## Funding, Acknowledgments, Declarations, etc

The authors thank Fitnat Yildiz and Kyle Floyd for kindly sharing the c-di-GMP reporter system and for information on fluorescent MSHA pilus labeling prior to publication. We are grateful to Ulrike Ruppert for her excellent technical support.

The study was funded by the Deutsche Forschungsgemeinschaft within the framework of the Priority Program SPP 1879 ‚Nucleotide second messenger signaling in bacteria’ (to KMT, GB and SR) and DFG GR1670/25-1 to PLG.

## Notes

### Competing Interest Statement

The authors have declared no competing interest.

### Summary of Updates

Line numbers have been added.

## References

1. U. Jenal, A. Reinders, C. Lori, Cyclic di-GMP: second messenger extraordinaire. Nat. Rev. Microbiol. 15, 271–284 (2017).

2. R. Hengge, Principles of c-di-GMP signalling in bacteria. Nat. Rev. Microbiol. 7, 263–273 (2009).

3. U. Römling, M. Y. Galperin, M. Gomelsky, Cyclic di-GMP: the first 25 years of a universal bacterial second messenger. Microbiol. Mol. Biol. Rev. MMBR 77, 1–52 (2013).

4. T. Schirmer, U. Jenal, Structural and mechanistic determinants of c-di-GMP signalling. Nat. Rev. Microbiol. 7, 724–735 (2009).

5. M. Valentini, A. Filloux, Multiple Roles of c-di-GMP signaling in bacterial pathogenesis. Annu. Rev. Microbiol. 73, 387–406 (2019).

6. S. Kunz, P. L. Graumann, Spatial organization enhances versatility and specificity in cyclic di-GMP signaling. Biol. Chem. 401, 1323–1334 (2020).

7. R. Hengge, High-specificity local and global c-di-GMP signaling. Trends Microbiol. (2021) https://doi.org/10.1016/j.tim.2021.02.003.

8. C. Lori, et al., Cyclic di-GMP acts as a cell cycle oscillator to drive chromosome replication. Nature 523, 236–239 (2015).

9. S. Abel, et al., Bi-modal distribution of the second messenger c-di-GMP controls cell fate and asymmetry during the Caulobacter cell cycle. PLoS Genet. 9, e1003744 (2013).

10. S. Abel, et al., Regulatory cohesion of cell cycle and cell differentiation through interlinked phosphorylation and second messenger networks. Mol. Cell 43, 550–560 (2011).

11. M. Christen, et al., Asymmetrical distribution of the second messenger c-di-GMP upon bacterial cell division. Science 328, 1295–1297 (2010).

12. B. R. Kulasekara, et al., c-di-GMP heterogeneity is generated by the chemotaxis machinery to regulate flagellar motility. eLife 2, e01402 (2013).

13. B.-J. Laventie, et al., A surface-induced asymmetric program promotes tissue colonization by *Pseudomonas aeruginosa*. Cell Host Microbe 25, 140–152.e6 (2019).

14. L. Chao, S. Rakshe, M. Leff, A. M. Spormann, PdeB, a cyclic Di-GMP-specific phosphodiesterase that regulates *Shewanella oneidensis* MR-1 motility and biofilm formation. J. Bacteriol. 195, 3827–3833 (2013).

15. F. M. Rossmann, et al., The GGDEF domain of the phosphodiesterase PdeB in *Shewanella putrefaciens* mediates recruitment by the polar landmark protein HubP. J. Bacteriol. 201 (2019).

16. Y. Yamaichi, et al., A multidomain hub anchors the chromosome segregation and chemotactic machinery to the bacterial pole. Genes Dev. 26, 2348–2360 (2012).

17. F. Rossmann, et al., The role of FlhF and HubP as polar landmark proteins in *Shewanella putrefaciens* CN-32. Mol. Microbiol. 98, 727–742 (2015).

18. N. Takekawa, S. Kwon, N. Nishioka, S. Kojima, M. Homma, HubP, a polar landmark protein, regulates flagellar number by assisting in the proper polar localization of FlhG in *Vibrio alginolyticus*. J. Bacteriol. 198, 3091–3098 (2016).

19. S. Inaba, T. Nishigaki, N. Takekawa, S. Kojima, M. Homma, Localization and domain characterization of the SflA regulator of flagellar formation in *Vibrio alginolyticus*. Genes Cells Devoted Mol. Cell. Mech. 22, 619–627 (2017).

20. S. Brenzinger, et al., ZomB is essential for flagellar motor reversals in *Shewanella putrefaciens* and *Vibrio parahaemolyticus*. Mol. Microbiol. 109, 694–709 (2018).

21. S. Bense, et al., Spatiotemporal control of FlgZ activity impacts *Pseudomonas aeruginosa* flagellar motility. Mol. Microbiol. 111, 1544–1557 (2019).

22. S. Park, et al., Polar landmark protein HubP recruits flagella assembly protein FapA under glucose limitation in *Vibrio vulnificus*. Mol. Microbiol. 112, 266–279 (2019).

23. M. Schniederberend, et al., Modulation of flagellar rotation in surface-attached bacteria: A pathway for rapid surface-sensing after flagellar attachment. PLoS Pathog. 15, e1008149 (2019).

24. A. B. T. Semmler, C. B. Whitchurch, A. J. Leech, J. S. Mattick, Identification of a novel gene, *fimV*, involved in twitching motility in *Pseudomonas aeruginosa*. Microbiol. Read. Engl. 146 (Pt 6), 1321–1332 (2000).

25. H. Wehbi, et al., The peptidoglycan-binding protein FimV promotes assembly of the *Pseudomonas aeruginosa* type IV pilus secretin. J. Bacteriol. 193, 540–550 (2011).

26. R. N. C. Buensuceso, et al., The conserved tetratricopeptide repeat-containing C-terminal domain of *Pseudomonas aeruginosa* FimV is required for its cyclic AMP-dependent and - independent functions. J. Bacteriol. 198, 2263–2274 (2016).

27. Y. F. Inclan, et al., A scaffold protein connects type IV pili with the Chp chemosensory system to mediate activation of virulence signaling in *Pseudomonas aeruginosa*. Mol. Microbiol. 101, 590–605 (2016).

28. R. N. C. Buensuceso, et al., Cyclic AMP-independent control of twitching motility in *Pseudomonas aeruginosa*. J. Bacteriol. 199 (2017).

29. T. Carter, et al., The type IVa pilus machinery is recruited to sites of future cell division. mBio 8 (2017).

30. T. Schirmer, c-di-GMP synthesis: Structural aspects of evolution, catalysis and regulation. J. Mol. Biol. 428, 3683–3701 (2016).

31. K. M. Thormann, et al., Control of formation and cellular detachment from *Shewanella oneidensis* MR-1 biofilms by cyclic di-GMP. J. Bacteriol. 188, 2681–2691 (2006).

32. H. Zhou, et al., Characterization of a natural triple-tandem c-di-GMP riboswitch and application of the riboswitch-based dual-fluorescence reporter. Sci. Rep. 6, 20871 (2016).

33. D. Zamorano-Sánchez, et al., Functional specialization in *Vibrio cholerae* diguanylate cyclases: Distinct modes of motility suppression and c-di-GMP production. mBio 10, e00670–19 (2019).

34. S. Theunissen, et al., The 285 kDa Bap/RTX hybrid cell surface protein (SO4317) of *Shewanella oneidensis* MR-1 is a key mediator of biofilm formation. Res. Microbiol. 161, 144–152 (2010).

35. C. Wu, et al., Oxygen promotes biofilm formation of *Shewanella putrefaciens* CN32 through a diguanylate cyclase and an adhesin. Sci. Rep. 3, 1945 (2013).

36. G. Zhou, J. Yuan, H. Gao, Regulation of biofilm formation by BpfA, BpfD, and BpfG in *Shewanella oneidensis*. Front. Microbiol. 6, 790 (2015).

37. K. M. Thormann, R. M. Saville, S. Shukla, D. A. Pelletier, A. M. Spormann, Initial Phases of biofilm formation in *Shewanella oneidensis* MR-1. J. Bacteriol. 186, 8096–8104 (2004).

38. R. M. Saville, N. Dieckmann, A. M. Spormann, Spatiotemporal activity of the *mshA* gene system in *Shewanella oneidensis* MR-1 biofilms. FEMS Microbiol. Lett. 308, 76–83 (2010).

39. C. J. Jones, et al., c-di-GMP regulates motile to sessile transition by modulating MshA pili biogenesis and near-surface motility behavior in *Vibrio cholerae*. PLoS Pathog. 11, e1005068 (2015).

40. Y.-C. Wang, et al., Nucleotide binding by the widespread high-affinity cyclic di-GMP receptor MshEN domain. Nat. Commun. 7, 12481 (2016).

41. K. A. Floyd, et al., c-di-GMP modulates type IV MSHA pilus retraction and surface attachment in *Vibrio cholerae*. Nat. Commun. 11, 1549 (2020).

42. A. S. N. Seshasayee, G. M. Fraser, N. M. Luscombe, Comparative genomics of cyclic-di-GMP signalling in bacteria: post-translational regulation and catalytic activity. Nucleic Acids Res. 38, 5970–5981 (2010).

43. R. B. R. Ferreira, L. C. M. Antunes, E. P. Greenberg, L. L. McCarter, *Vibrio parahaemolyticus* ScrC modulates cyclic dimeric GMP regulation of gene expression relevant to growth on surfaces. J. Bacteriol. 190, 851–860 (2008).

44. M. Tarutina, D. A. Ryjenkov, M. Gomelsky, An unorthodox bacteriophytochrome from *Rhodobacter sphaeroides* involved in turnover of the second messenger c-di-GMP*. J. Biol. Chem. 281, 34751–34758 (2006).

45. C. W. Phippen, et al., Formation and dimerization of the phosphodiesterase active site of the *Pseudomonas aeruginosa* MorA, a bi-functional c-di-GMP regulator. FEBS Lett. 588, 4631–4636 (2014).

46. M. Christen, B. Christen, M. Folcher, A. Schauerte, U. Jenal, Identification and characterization of a cyclic di-GMP-specific phosphodiesterase and its allosteric control by GTP*. J. Biol. Chem. 280, 30829–30837 (2005).

47. S. An, J. Wu, L.-H. Zhang, Modulation of *Pseudomonas aeruginosa* biofilm dispersal by a cyclic-di-GMP phosphodiesterase with a putative hypoxia-sensing domain. Appl. Environ. Microbiol. 76, 8160–8173 (2010).

48. C. Liu, et al., Insights into biofilm dispersal regulation from the crystal structure of the PAS-GGDEF-EAL region of RbdA from *Pseudomonas aeruginosa*. J. Bacteriol. 200 (2018).

49. F. Mantoni, et al., Insights into the GTP-dependent allosteric control of c-di-GMP hydrolysis from the crystal structure of PA0575 protein from *Pseudomonas aeruginosa*. FEBS J. 285, 3815–3834 (2018).

50. G. G. Nicastro, et al., c-di-GMP-related phenotypes are modulated by the interaction between a diguanylate cyclase and a polar hub protein. Sci. Rep. 10, 3077 (2020).

51. S. Bubendorfer, et al., Specificity of motor components in the dual flagellar system *of Shewanella putrefaciens* CN-32. Mol. Microbiol. 83, 335–350 (2012).

52. S. Bubendorfer, M. Koltai, F. Rossmann, V. Sourjik, K. M. Thormann, Secondary bacterial flagellar system improves bacterial spreading by increasing the directional persistence of swimming. Proc. Natl. Acad. Sci. U. S. A. 111, 11485–11490 (2014).

53. Y.-Y. Cheng, et al., FlrA represses transcription of the biofilm-associated *bpfA* operon in *Shewanella putrefaciens*. Appl. Environ. Microbiol. 83 (2017).

54. A. Pecina, et al., The stand-alone PilZ-domain protein MotL specifically regulates the activity of the secondary lateral flagellar system in *Shewanella putrefaciens*. Front. Microbiol. 12, 668892 (2021).

55. J. F. Lebov, B. J. M. Bohannan, Msh pilus mutations increase the ability of a free-living bacterium to colonize a piscine host. Genes 12 (2021).

56. K. G. Roelofs, et al., Systematic identification of cyclic-di-GMP binding proteins in *Vibrio cholerae* reveals a novel class of cyclic-di-GMP-binding ATPases associated with type II secretion systems. PLoS Pathog. 11, e1005232 (2015).

57. A. J. Collins, T. J. Smith, H. Sondermann, G. A. O’Toole, From Input to Output: The Lap/c-di-GMP biofilm regulatory circuit. Annu. Rev. Microbiol. 74, 607–631 (2020).

58. H. H. Hau, J. A. Gralnick, Ecology and biotechnology of the genus *Shewanella*. Annu. Rev. Microbiol. 61, 237–258 (2007).

59. E. C. Stuffle, M. S. Johnson, K. J. Watts, PAS domains in bacterial signal transduction. Curr. Opin. Microbiol. 61, 8–15 (2021).

60. D. G. Gibson, et al., Enzymatic assembly of DNA molecules up to several hundred kilobases. Nat. Methods 6, 343–345 (2009).

61. J. Lassak, A.-L. Henche, L. Binnenkade, K. M. Thormann, ArcS, the cognate sensor kinase in an atypical Arc system of *Shewanella oneidensis* MR-1. Appl. Environ. Microbiol. 76, 3263–3274 (2010).

62. C. A. Schneider, W. S. Rasband, K. W. Eliceiri, NIH Image to ImageJ: 25 years of image analysis. Nat. Methods 9, 671–675 (2012).

63. C. K. Ellison, T. N. Dalia, A. B. Dalia, Y. V. Brun, Real-time microscopy and physical perturbation of bacterial pili using maleimide-conjugated molecules. Nat. Protoc. 14, 1803–1819 (2019).

64. M. J. Kühn, F. K. Schmidt, B. Eckhardt, K. M. Thormann, Bacteria exploit a polymorphic instability of the flagellar filament to escape from traps. Proc. Natl. Acad. Sci. U. S. A. 114, 6340–6345 (2017).

65. R. Hartmann, M. C. F. van Teeseling, M. Thanbichler, K. Drescher, BacStalk: A comprehensive and interactive image analysis software tool for bacterial cell biology. Mol. Microbiol. 114, 140–150 (2020).

66. J. Schindelin, et al., Fiji: an open-source platform for biological-image analysis. Nat. Methods 9, 676–682 (2012).

67. A. Paintdakhi, et al., Oufti: an integrated software package for high-accuracy, high-throughput quantitative microscopy analysis. Mol. Microbiol. 99, 767–777 (2016).

68. K. Jaqaman, et al., Robust single-particle tracking in live-cell time-lapse sequences. Nat. Methods 5, 695–702 (2008).

69. W. Steinchen, et al., Catalytic mechanism and allosteric regulation of an oligomeric (p)ppGpp synthetase by an alarmone. Proc. Natl. Acad. Sci. U. S. A. 112, 13348–13353 (2015).

70. T. E. Wales, K. E. Fadgen, G. C. Gerhardt, J. R. Engen, High-speed and high-resolution UPLC separation at zero degrees Celsius. Anal. Chem. 80, 6815–6820 (2008).

71. S. J. Geromanos, et al., The detection, correlation, and comparison of peptide precursor and product ions from data independent LC-MS with data dependant LC-MS/MS. Proteomics 9, 1683–1695 (2009).

72. G.-Z. Li, et al., Database searching and accounting of multiplexed precursor and product ion spectra from the data independent analysis of simple and complex peptide mixtures. Proteomics 9, 1696–1719 (2009).

73. M. Osorio-Valeriano, et al., ParB-type DNA segregation proteins are CTP-dependent molecular switches. Cell 179, 1512–1524.e15 (2019).

74. M. Götze, et al., Automated assignment of MS/MS cleavable cross-links in protein 3D-structure analysis. J. Am. Soc. Mass Spectrom. 26, 83–97 (2015).

75. F. Sievers, et al., Fast, scalable generation of high-quality protein multiple sequence alignments using Clustal Omega. Mol. Syst. Biol. 7, 539 (2011).

76. A. M. Waterhouse, J. B. Procter, D. M. A. Martin, M. Clamp, G. J. Barton, Jalview Version 2--a multiple sequence alignment editor and analysis workbench. Bioinforma. Oxf. Engl. 25, 1189–1191 (2009).

77. M. C. F. Thomsen, M. Nielsen, Seq2Logo: a method for construction and visualization of amino acid binding motifs and sequence profiles including sequence weighting, pseudo counts and two-sided representation of amino acid enrichment and depletion. Nucleic Acids Res. 40, W281–W287 (2012).

78. L. A. Kelley, S. Mezulis, C. M. Yates, M. N. Wass, M. J. E. Sternberg, The Phyre2 web portal for protein modeling, prediction and analysis. Nat. Protoc. 10, 845–858 (2015).

79. H. Bähre, V. Kaever, Identification and quantification of cyclic di-Guanosine monophosphate and its linear metabolites by Reversed-Phase LC-MS/MS. Methods Mol. Biol. Clifton NJ 1657, 45–58 (2017).

