## Supplemental Tables 1-3 and Supplemental Figures 1-8 for "A GGDEF domain serves as a spatial on-switch for a phosphodiesterase by direct interaction with a polar landmark protein"

### - SUPPLEMENTAL INFORMATION -

Tim Rick[1], Vanessa Kreiling[1], Alexander Höing[2], Svenja Fiedler[3,4], Timo Glatter[5], Wieland Steinchen[3,4], Georg Hochberg[6], Heike Bähre[7], Roland Seifert[8], Gert Bange[3,4], Shirley K. Knauer[2], Peter L. Graumann[3,4], Kai M. Thormann[1]

[1]Justus-Liebig-Universität, Department of Microbiology and Molecular Biology, 35392 Giessen, Germany

[2]Department of Molecular Biology II, Centre for Medical Biotechnology (ZMB), University of Duisburg-Essen, 45141 Essen, Germany

[3]LOEWE Center for Synthetic Microbiology, SYNMIKRO, 35043 Marburg, Germany

[4]Philipps-Universität Marburg, Department of Chemistry, 35043 Marburg, Marburg, Germany

[5]Max Planck Institute for Terrestrial Microbiology, Facility for Mass Spectrometry and Proteomics 35043 Marburg, Germany

[6]Max Planck Institute for Terrestrial Microbiology, 35043 Marburg, Germany

[7]Medizinische Hochschule Hannover, ZFA Metabolomics, 30625 Hannover, Germany

### Contents: Supplemental Tables 1 – 3

### Supplemental Figures 1 – 9

### Supplemental Table 1: Bacterial strains that were used in this study

| Strain | Genotype | Purpose | Reference |
|---|---|---|---|
| ***Escherichia coli*** | | | |
| **DH5α λpir** | φ80d*lacZ* ΔM15 Δ(*lacZYA-argF*)U169 *recA*$_1$ *hsdR17 deoR thi-l supE44 gyrA96 relA*$_1$*/λpir* | cloning strain | (1) |
| **WM3064** | *thrB1004 pro thi rpsL hsdS lacZ* ΔM15 RP4-1360 Δ(*araBAD*) 567Δ*dapA* 1341::[*erm pir*(wt)] | conjugation strain for *Shewanella* | W. Metcalf, University of Illinois, Urbana-Champaign |
| **BL21(DE3)** | *fhuA2* [*lon*] *ompT gal* (λ *DE3*) [*dcm*] Δ*hsdS* λ *DE3* = λ *sBamHIo* Δ*EcoRI-B int*::(*lacI*::*PlacUV5*::*T7 gene1*) *i21* Δ*nin5* | protein overproduction strain | New England Biolabs (NEB) |
| ***Shewanella putrefaciens* CN-32** | | | |
| **S757** | CN-32 wt | wild type | (2) |
| **S2576** | Δ*flaAB*$_2$ | markerless in-frame deletion of the lateral flagellins (*sputcn32_3455-3456*) | (3) |
| **S3297** | Δ*pdeB* | markerless in-frame deletion of the gene *sputcn32_3405 (pdeB)* | (4) |
| **S4234** | *pdeB-sfgfp* | markerless in-frame fusion of *pdeB* with sf*gfp* | (4) |
| **S4237** | *pdeB*-E637A-*sfgfp* | functional markerless substitution of the EIL motif to AIL (residue 637) in the background of *pdeB-sfgfp* | (4) |
| **S4357** | Δ*flaAB*$_2$ Δ*pdeB* | markerless in-frame deletion of the gene *pdeB* (*sputcn32_3405*) in a background with deleted lateral flagellins | (4) |
| **S6452** | *pdeB-gfp* V522G V523G Q524G | markerless in-frame fusion of *pdeB* with sf*gfp* substitution of Q524G | this study |
| **S6453** | *pdeB-gfp* K490G V491G M492G Q593G | markerless in-frame fusion of *pdeB* with sf*gfp* substitution of M492G Q593G | this study |
| **S6454** | *pdeB-GFP* R557G A558G P559G Y560G | markerless in-frame fusion of *pdeB* with *sfgfp* substitution of A558G P559G Y560G | this study |
| **S6496** | Δ*flaAB*$_2$ Δ*mshE* | markerless in-frame deletion of the gene *mshE (sputcn32_0563)* in a background with deleted lateral flagellins | this study |
| **S6497** | Δ*flaAB*$_2$ Δ*pdeB* Δ*mshE* | markerless in-frame deletion of the gene *mshE (sputcn32_0563)* in a background with deleted lateral flagellins and deletion of the gene *pdeB* (*sputcn32_3405*) | this study |
| **S6527** | *pdeB-gfp* K490D Q493A | markerless in-frame fusion of *pdeB* with sf*gfp* substitution of K490D Q493A | this study |
| **S6528** | *pdeB-gfp* Q524A K527D Q528A | markerless in-frame fusion of *pdeB* with sf*gfp* substitution of Q524A K527D Q528A | this study |
| **S6683** | Δ*flaAB*$_2$ *mshA* S68C | functional substitution of *mshA* S68C in a background with deleted lateral flagellins | this study |
| **S6684** | Δ*flaAB*$_2$ Δ*pdeB mshA* S68C | functional substitution of *mshA* S68C in a background with deleted lateral flagellins and deletion of the gene *pdeB* (*sputcn32_3405*) | this study |

| | | | |
|---|---|---|---|
| **S6688** | Δ*flaAB*$_2$ Δ*aggA* | markerless in-frame deletion of the gene *aggA (sputcn32_3594)* in a background with deleted lateral flagellins | this study |
| **S6689** | Δ*flaAB*$_2$ Δ*pdeB* Δ*aggA* | markerless in-frame deletion of the gene *aggA (sputcn32_3594)* in a background with deleted lateral flagellins and deletion of the gene *pdeB (sputcn32_3405)* | this study |
| **S6690** | Δ*flaAB*$_2$ Δ*pdeB* Δ*mshE* Δ*aggA* | markerless in-frame deletion of the gene *aggA (sputcn32_3594)* in a background with deleted lateral flagellins, and deletion of the genes *pdeB (sputcn32_3405)* and *mshE (sputcn32_0563)* | this study |
| **S6691** | Δ*flaAB*$_2$ Δ*pilB* | markerless in-frame deletion of the gene *pilB (sputcn32_3423)* in a background with deleted lateral flagellins | this study |
| **S6692** | Δ*flaAB*$_2$ Δ*pdeB* Δ*pilB* | markerless in-frame deletion of the gene *pilB (sputcn32_3423)* in a background with deleted lateral flagellins and deletion of the gene *pdeB (sputcn32_3405)* | this study |
| **S6693** | Δ*flaAB*$_2$ Δ*pdeB* Δ*mshE* Δ*pilB* | markerless in-frame deletion of the gene *pilB (sputcn32_3423)* in a background with deleted lateral flagellins, and deletion of the genes *pdeB (sputcn32_3405)* and *mshE (sputcn32_0563)* | this study |
| **S6729** | *pdeB-gfp* K527E Q528S | markerless in-frame fusion of *pdeB* with sf*gfp* substitution of K527E Q528S | this study |
| **S7024** | *pdeB-mvenus* | markerless in-frame fusion of *pdeB* with *mvenus* | this study |
| **S7025** | *pdeB-mvenus* D508A E509A | markerless in-frame fusion of *pdeB* D508A E509A with *mvenus* | this study |
| **S7026** | *pdeB-mvenus* E637A | markerless in-frame fusion of *pdeB* E637A with *mvenus* | this study |
| **S7243** | *pdeB-gfp* K527S Q528S | markerless in-frame fusion of *pdeB* with sf*gfp* substitution of K527S Q528S | this study |
| **S7243** | *pdeB-gfp* Q524S Q528S | markerless in-frame fusion of *pdeB* with sf*gfp* substitution of Q524S Q528S | this study |
| **S7244** | *pdeB-gfp* K527S | markerless in-frame fusion of *pdeB* with *sfgfp* substitution of K527S | this study |
| **S7245** | *pdeB-gfp* G497A | markerless in-frame fusion of *pdeB* with *sfgfp* substitution of G497A | this study |
| **S7246** | *pdeB-gfp Q499S* | markerless in-frame fusion of *pdeB* with *sfgfp* substitution of Q499S | this study |
| **S7247** | *pdeB-gfp* E500S | markerless in-frame fusion of *pdeB* with *sfgfp* substitution of E500S | this study |
| **S7444** | Δ*flaAB*$_2$ Δ*mshE* Δ*aggA* | markerless in-frame deletion of the gene *aggA (sputcn32_3594)* in a background with deleted lateral flagellins and deletion of the gene *mshE (sputcn32_0563)* | this study |
| **S7445** | Δ*flaAB*$_2$ Δ*mshE* Δ*pilB* | markerless in-frame deletion of the gene *pilB (sputcn32_3423)* in a background with deleted lateral flagellins and deletion of the gene *mshE (sputcn32_0563)* | this study |
| **S7504** | *pdeB* K527E Q528S | markerless in-frame substitution of *pdeB* K527E Q528S | this study |

| | | | |
|---|---|---|---|
| **S7505** | *pdeB* G497A | markerless in-frame substitution of *pdeB* G497A | this study |
| **S7506** | *pdeB* K578S | markerless in-frame substitution of *pdeB* K578S | this study |
| **S7507** | *pdeB-mvenus* K527E Q528S | markerless in-frame fusion of *pdeB*-K527E Q528S with *mvenus* | this study |
| **S7508** | *pdeB* K580S | markerless in-frame substitution of *pdeB* K580S | this study |
| **S7562** | *pdeB-gfp* K578S | markerless in-frame fusion of *pdeB* with *sfgfp* substitution of K578S | this study |
| **S7564** | *pdeB-gfp* K580S | markerless in-frame fusion of *pdeB* with *sfgfp* substitution of K580S | this study |
| **S7614** | pMMB-Gm-Bc3-5 AAV (hok-sok) | wildtype strain containing the c-di-GMP biosensor plasmid | this study |
| **S7616** | Δ*pdeB* pMMB-Gm-Bc3-5 AAV (hok-sok) | c-di-GMP biosensor plasmid in the background of deleted *pdeB* (*sputcn32_3405*) | this study |
| **S7653** | *pdeB* K527E Q528S pMMB-Gm-Bc3-5 AAV (hok-sok) | c-di-GMP biosensor plasmid in the background of *pdeB* K527E Q528S substitution | this study |
| **S7654** | *pdeB* G497A pMMB-Gm-Bc3-5 AAV (hok-sok) | c-di-GMP biosensor plasmid in the background of *pdeB* G497A substitution | this study |
| **S7655** | *pdeB* K578S pMMB-Gm-Bc3-5 AAV (hok-sok) | c-di-GMP biosensor plasmid in the background of *pdeB* K578S substitution | this study |
| **S7691** | *lapA*-GS-3xFLAG | functional markerless in-frame tag of 3xFLAG to the C-terminus of *lapA* via a flexible GS-linker | this study |
| **S7692** | Δ*pdeB lapA*-GS-3xFLAG | functional markerless in-frame tag of 3xFLAG to the C-terminus of *lapA* via a flexible GS-linker in the background of deleted *pdeB (sputcn32_3405)* | this study |
| **S7703** | *lapB*-GS-3xFLAG | functional markerless in-frame tag of 3xFLAG to the C-terminus of *lapB* via a flexible GS-linker | this study |
| **S7704** | Δ*pdeB lapB*-GS-3xFLAG | functional markerless in-frame tag of 3xFLAG to the C-terminus of *lapB* via a flexible GS-linker in the background of deleted *pdeB (sputcn32_3405)* | this study |
| ***Shewanella oneidensis* MR-1** | | | |
| **S79** | MR-1 wt | wildtype strain | (5) |
| **S7296** | Δ*pdeB* | Markerless in-frame deletion of *pdeB* of *S. oneidensis* MR-1 | this study |
| **S7294** | *pdeB-gfp* | Markerless in-frame fusion of *pdeB* with *sfgfp* in *S. oneidensis* MR-1 | this study |
| **S7423** | pMMB-Gm-Bc3-5 AAV (hok-sok) | MR-1 wildtype strain containing the c-di-GMP biosensor plasmid | this study |
| **S7425** | Δ*pdeB* pMMB-HS-Bc-3-5-AAV (hok-sok) | c-di-GMP biosensor plasmid in the background of deleted *pdeB* (*SO_0437*) | this study |

**Supplemental Table 2: Plasmids that were used in this study**

| Plasmid | Relevant genotype or phenotype | Reference |
|---|---|---|
| **pNPTS-138-R6KT** | *mob*RP4+ *ori*-R6K *sacB* β-galactosidase fragment alpha, suicide plasmid for in frame deletions/insertions in *Shewanella*, $Km^r$ | (6) |
| **pET-24c** | overproduction vector for His-tagged proteins | EMD Biosciences |
| **pBTOK** | pBBR1-MCS2 backbone (pBBR origin, $Km^r$ ); TetR, Promoter and multiple cloning site of pASK-IBA3plus and *E. coli* rrnB1 T1 and lambda phage T0 terminator. Overproduction plasmid, inducible with anhydrotetracycline | (7) |
| **pMMB-Gm-Bc3-5 AAV (hok-sok)** | pMMB67EH (Gm) backbone containing the c-di-GMP biosensor (turboRFP with an AAV tag) and the hok/sok region from pXB300. Used as c-di-GMP reporter. | Fitnat Yildiz, UCSC Santa Cruz, CA |
| **overexpression vectors** | | |
| **pET24c MBP-PdeB (MR-1) GGDEF-6xHis** | Vector used to express the GGDEF-domain of MR-1 PdeB (residues 417 - 585) with N-terminal MBP and C-terminal 6xHis translational fusion | this study |
| **pET24c MBP-PdeB (MR-1) GGDEF-6xHis K524S** | Vector used to express the GGDEF-domain of MR-1 PdeB (residues 417 - 585) with N-terminal MBP and C-terminal 6xHis translational fusion | this study |
| **pET24c MBP-PdeB (MR-1) GGDEF-6xHis Q525S** | Vector used to express the GGDEF-domain of MR-1 PdeB (residues 417 - 585) with N-terminal MBP and C-terminal 6xHis translational fusion | this study |
| **pET24c MBP-PdeB (MR-1) GGDEF-6xHis K524E Q525S** | Vector used to express the GGDEF-domain of MR-1 PdeB (residues 417 - 585) with N-terminal MBP and C-terminal 6xHis translational fusion | this study |
| **pET24c MBP-PdeB (MR-1) GGDEF-6xHis G494A** | Vector used to express the GGDEF-domain of MR-1 PdeB (residues 417 - 585) with N-terminal MBP and C-terminal 6xHis translational fusion | this study |
| **pET24c MBP-PdeB (MR-1) GGDEF-6xHis E497S** | Vector used to express the GGDEF-domain of MR-1 PdeB (residues 417 - 585) with N-terminal MBP and C-terminal 6xHis translational fusion | this study |
| **pET24c MBP-PdeB (MR-1) PAS-GGDEF-6xHis** | Vector used to express the PAS- and GGDEF-domain of MR-1 PdeB (residues 304 - 585) with N-terminal MBP and C-terminal 6xHis translational fusion | this study |
| **pET24c MBP-PdeB (MR-1) PAS-GGDEF-6xHis K524E Q525S** | Vector used to express the PAS- and GGDEF-domain of MR-1 PdeB (residues 304 - 585) with N-terminal MBP and C-terminal 6xHis translational fusion | this study |
| **pET24c (MR-1) FimV-Cdomain-6xHis** | Vector used to express the C-terminal domain of MR-1 FimV (residues 1000 - 1110) with C-terminal 6xHis translational fusion | this study |
| **pET24c 3xFLAG-(CN-32) HubP-FimV-Cdomain-6xHis** | Vector used to express the C-terminal domain of MR-1 FimV (residues 1000 - 1110) with C-terminal 6xHis translational fusion | this study |
| **pET24c *mshE*-6xHis** | Vector used to express MshE of CN-32 with C-terminal 6xHis translational fusion | this study |
| **pET24c *mshE*_Ndomain-6xHis** | Vector used to express the N-terminal domain of CN-32 MshE (residues 2 - 145) with C-terminal 6xHis translational fusion | this study |
| **pET24c *pilB*_Ndomain-6xHis** | Vector used to express the N-terminal domain of CN-32 PilB (residues 2 - 145) with C-terminal 6xHis translational fusion | this study |
| **pBTOK *dgcA*-6xHis** | Vector for ectopical expression of *dgcA* (*E. coli*) in *S. putrefaciens* CN-32 with C-terminal 6xHis | this study |
| **pBTOK *dgcA*-6xHis D216E** | Vector for ectopical expression of *dgcA* (*E. coli*) in *S. putrefaciens* CN-32 with C-terminal 6xHis and D216E | this study |

| | | |
|---|---|---|
| **pBTOK *dgcA*-6xHis E276K** | Vector for ectopical expression of *dgcA* (*E. coli*) in *S. putrefaciens* CN-32 with C-terminal 6xHis and E276K | this study |
| **In-frame insertion vectors** | | |
| **pNPTS CN-32 *pdeB-gfp* K527S** | Suicide vector for markerless in-frame insertion of *pdeB-gfp* of *S. putrefaciens* CN-32 with K527S | this study |
| **pNPTS CN-32 *pdeB-gfp* K527S Q528S** | Suicide vector for markerless in-frame insertion of *pdeB-gfp* of *S. putrefaciens* CN-32 with K527S Q528S | this study |
| **pNPTS CN-32 *pdeB-gfp* K527D** | Suicide vector for markerless in-frame insertion of *pdeB-gfp* of *S. putrefaciens* CN-32 with K527D | this study |
| **pNPTS CN-32 *pdeB-gfp* K527D Q528S** | Suicide vector for markerless in-frame insertion of *pdeB-gfp* of *S. putrefaciens* CN-32 with K527D Q528S | this study |
| **pNPTS CN-32 *pdeB-gfp* Q524A K527D Q528A** | Suicide vector for markerless in-frame insertion of *pdeB-gfp* of *S. putrefaciens* CN-32 with Q524A K527D Q528A | this study |
| **pNPTS CN-32 *pdeB-gfp* G497A** | Suicide vector for markerless in-frame insertion of *pdeB-gfp* of *S. putrefaciens* CN-32 with G497A | this study |
| **pNPTS CN-32 *pdeB-gfp* Q499S** | Suicide vector for markerless in-frame insertion of *pdeB-gfp* of *S. putrefaciens* CN-32 with Q499S | this study |
| **pNPTS CN-32 *pdeB-gfp* E500S** | Suicide vector for markerless in-frame insertion of *pdeB-gfp* of *S. putrefaciens* CN-32 with E500S | this study |
| **pNPTS CN-32 *pdeB-gfp* Q528S** | Suicide vector for markerless in-frame insertion of *pdeB-gfp* of *S. putrefaciens* CN-32 with Q528S | this study |
| **pNPTS CN-32 *pdeB-gfp* Q524S** | Suicide vector for markerless in-frame insertion of *pdeB-gfp* of *S. putrefaciens* CN-32 with Q524S | this study |
| **pNPTS CN-32 *pdeB-gfp* Q524S Q528S** | Suicide vector for markerless in-frame insertion of *pdeB-gfp* of *S. putrefaciens* CN-32 with Q524S Q528S | this study |
| **pNPTS CN-32 *pdeB-gfp* K527E Q528S** | Suicide vector for markerless in-frame insertion of *pdeB-gfp* of *S. putrefaciens* CN-32 with K527E Q528S | this study |
| **pNPTS CN-32 *pdeB-gfp* K490D Q493A** | Suicide vector for markerless in-frame insertion of *pdeB-gfp* of *S. putrefaciens* CN-32 with K490D Q493A | this study |
| **pNPTS CN-32 *pdeB-gfp* R557G A558G P559G Y560G** | Suicide vector for markerless in-frame insertion of *pdeB-gfp* of *S. putrefaciens* CN-32 with R557G A558G P559G Y560G | this study |
| **pNPTS CN-32 *pdeB-gfp* V522G V523G Q524G** | Suicide vector for markerless in-frame insertion of *pdeB-gfp* of *S. putrefaciens* CN-32 with V522G V523G Q524G | this study |
| **pNPTS CN-32 *pdeB-gfp* K490G V491G M492G Q593G** | Suicide vector for markerless in-frame insertion of *pdeB-gfp* of *S. putrefaciens* CN-32 with K490G V491G M492G Q593G | this study |
| **pNPTS CN-32 *pdeB-gfp* K578S** | Suicide vector for markerless in-frame insertion of *pdeB-gfp* of *S. putrefaciens* CN-32 with K578S | this study |
| **pNPTS CN-32 *pdeB-gfp* K580S** | Suicide vector for markerless in-frame insertion of *pdeB-gfp* of *S. putrefaciens* CN-32 with K580S | this study |
| **pNPTS CN-32 *pdeB* G497A** | Suicide vector for markerless in-frame insertion of *pdeB* of *S. putrefaciens* CN-32 with G497A | this study |
| **pNPTS CN-32 *pdeB* K527E Q528S** | Suicide vector for markerless in-frame insertion of *pdeB* of *S. putrefaciens* CN-32 with K527E Q528S | this study |
| **pNPTS CN-32 *pdeB* K578S** | Suicide vector for markerless in-frame insertion of *pdeB* of *S. putrefaciens* CN-32 with K578S | this study |
| **pNPTS CN-32 *pdeB* K580S** | Suicide vector for markerless in-frame insertion of *pdeB* of *S. putrefaciens* CN-32 with K580S | this study |
| **pNPTS CN-32 *pdeB-venus*** | Suicide vector for markerless in-frame insertion of *pdeB-mvenus* of *S. putrefaciens* CN-32 | this study |
| **pNPTS CN-32 *pdeB-venus* D508A E509A** | Suicide vector for markerless in-frame insertion of *pdeB-mvenus* of *S. putrefaciens* CN-32 with 508A E509A | this study |
| **pNPTS CN-32 *pdeB-venus* E637A** | Suicide vector for markerless in-frame insertion of *pdeB-mvenus* of *S. putrefaciens* CN-32 with E637A | this study |
| **pNPTS CN-32 *lapA*-GS-3xFLAG** | Suicide vector for markerless in-frame insertion of 3xFLAG to the C-terminus of *lapA* via a flexible GS-linker | this study |
| **pNPTS CN-32 *aggC*-GS-3xFLAG** | Suicide vector for markerless in-frame insertion of 3xFLAG to the C-terminus of *aggC* via a flexible GS-linker | this study |
| **pNPTS CN-32 *mshA* S68C** | Suicide vector for markerless in-frame insertion of *mshA* of *S. putrefaciens* CN-32 with S68C | this study |

| | | |
|---|---|---|
| **pNPTS MR-1 *pdeB-gfp*** | Suicide vector for markerless in-frame insertion of *pdeB-gfp* of *S. oneidensis* MR-1 | this study |
| **In-frame deletion vectors** | | |
| **pNPTS CN-32 Δ*mshE*** | Suicide vector for markerless in-frame deletion of *mshE* of *S. putrefaciens* CN-32 | this study |
| **pNPTS CN-32 Δ*aggA*** | Suicide vector for markerless in-frame deletion of *aggA* of *S. putrefaciens* CN-32 | this study |
| **pNPTS CN-32 Δ*pilB*** | Suicide vector for markerless in-frame deletion of *pilB* of *S. putrefaciens* CN-32 | this study |
| **pNPTS MR-1 Δ*pdeB*** | Suicide vector for markerless in-frame deletion of *pdeB* of *S. oneidensis* MR-1 | this study |

**Supplemental Table 3: Oligonucleotides/primers that were used in this study**

| Plasmid | Primer | Sequence |
|---|---|---|
| **pET24c MBP-PdeB (MR-1) GGDEF-6xHis** | TR258 MBP fw | TTAACTTTAAGAAGGAGATATACAATGAAAATAGAAGAAGGTAAACTGGTAATCTGG |
| | TR259 MBP rv | GCTGCCCCCGAGGTTGTTGTTATTGTTATTGT |
| | TR260 | AATAACAACAACCTCGGGGGCAGCGAAGAACTTCTTAAGCATCAGCTAC |
| | TR257 | GTGGTGGTGGTGGTGGTGCTCAATGGTGATGGTGATGGTGGTAAATGTGGATTTGGTTGGTGC |
| **pET24c MBP-PdeB (MR-1) GGDEF-6xHis K524S** | TR510 MBP ol plas fw | TTAACTTTAAGAAGGAGATATACAATGAAAATAGAAGAAGGTAAACTGGTAATCTGG |
| | TR511 So KtoS fw | CAATAATTTGGCTCAGCAACTGCGCCACAGC |
| | TR512 So KtoS rv | GTTGCTGAGCCAAATTATTGCTCAAGTATCGCTGC |
| | TR513 soGGDEF ol plas rv | GTGGTGGTGGTGGTGGTGCTCAATGGTGATGGTGATGGTGGTAAATGTGGATTTGGTTGGTGC |
| **pET24c MBP-PdeB (MR-1) GGDEF-6xHis Q525S** | TR510 MBP ol plas fw | TTAACTTTAAGAAGGAGATATACAATGAAAATAGAAGAAGGTAAACTGGTAATCTGG |
| | TR514 So QtoS fw | GAGCAATAATGCTCTTCAGCAACTGCGCCACAGC |
| | TR515 So QtoS rv | GCTGAAGAGCATTATTGCTCAAGTATCGCTGC |
| | TR513 soGGDEF ol plas rv | GTGGTGGTGGTGGTGGTGCTCAATGGTGATGGTGATGGTGGTAAATGTGGATTTGGTTGGTGC |
| **pET24c MBP-PdeB (MR-1) GGDEF-6xHis K524E Q525S** | TR510 MBP ol plas fw | TTAACTTTAAGAAGGAGATATACAATGAAAATAGAAGAAGGTAAACTGGTAATCTGG |
| | TR593 SO KQ to ES rv | TAATGCTTTCCAGCAACTGCGCCACAGCTAA |
| | TR594 SO KQ to ES fw | GCAGTTGCTGGAAAGCATTATTGCTCAAGTATCGCTGCAAGTG |
| | TR513 soGGDEF ol plas rv | GTGGTGGTGGTGGTGGTGCTCAATGGTGATGGTGATGGTGGTAAATGTGGATTTGGTTGGTGC |
| **pET24c MBP-PdeB (MR-1) GGDEF-6xHis G494A** | TR510 MBP ol plas fw | TTAACTTTAAGAAGGAGATATACAATGAAAATAGAAGAAGGTAAACTGGTAATCTGG |
| | TR544 G494A fw | ATTCCTGTGGCGCAAGACATGACTGAATCGCCCTAG |
| | TR555 G494A rv | ATGTCTTGCGCCACAGGAATTATTAGCCCGCA |
| | TR513 soGGDEF ol plas rv | GTGGTGGTGGTGGTGGTGCTCAATGGTGATGGTGATGGTGGTAAATGTGGATTTGGTTGGTGC |
| **pET24c MBP-PdeB (MR-1) GGDEF-6xHis E497S** | TR510 MBP ol plas fw | TTAACTTTAAGAAGGAGATATACAATGAAAATAGAAGAAGGTAAACTGGTAATCTGG |
| | TR558 E497S fw | GGGCTAATAACGACTGTGGCCCAAGACATGACT |

| | | |
|---|---|---|
| | TR559 E497S rv | GCCACAGTCGTTATTAGCCCGCATAGGAGGTG |
| | TR513 soGGDEF ol plas rv | GTGGTGGTGGTGGTGGTGCTCAATGGTGATGGTGATGGTGGTAAATGTGGATTTGGTTGGTGC |
| **pET24c MBP-PdeB (MR-1) PAS-GGDEF-6xHis** | TR588 MBP fw SO | TTAACTTTAAGAAGGAGATATACAATGAAAATAGAAGAAGGTAAACTGGTAATCTGG |
| | TR589 MBP rv OL SO | TACCGCGCTCCCCGAGGTTGTTGTTATTGTTATTGT |
| | TR590 pet SO PAS fw | CAACCTCGGGGAGCGCGGTAAAATAACCTTAGA |
| | TR591 pet rv | GTGGTGGTGGTGGTGGTGCTCAATGGTGATGGTGATGGTGGTAAATGTGGATTTGGTTGGTGC |
| **pET24c mshE-6xHis** | TR258 MshE OW fw | TTAACTTTAAGAAGGAGATATACAATGAAACCCAGATTAAAGATGCGTTT |
| | TR259 MshE OW rv | GTGGTGGTGGTGGTGGTGCTCAATGGTGATGGTGATGGTGCGCCTCAACGCCTTGTTGG |
| **pET24c mshE_Ndomain-6xHis** | TR372 | TTAACTTTAAGAAGGAGATATACAATGCACCATCACCATCACCATAAACCCAGATTAAAGATGCGTTTGG |
| | TR383 | GTGGTGGTGGTGGTGGTGCCTAACGACGATAAAGATTATCAAAGGCC |
| **pET24c pilB_Ndomain-6xHis** | TR374 | TTAACTTTAAGAAGGAGATATACAATGCACCATCACCATCACCATATGCCAACCACTGGTCTTCATTTA |
| | TR384 | GTGGTGGTGGTGGTGGTGCCTATTCAAGGATTTTTTCAAGGGCTTTAG |
| **pNPTS CN-32 Δ*aggA*** | TR377 | GCGAATTCGTGGATCCAGATTGAAATCAGCCCTAGACGAAGC |
| | TR378 | TGTTAGTTCCTACTAAAGTATTCATTGCAAACCTCC |
| | TR379 | TACTTTAGTAGGAACTAACAAATGAAAACCGTAATC |
| | TR380 | GCCAAGCTTCTCTGCAGGATGGAGTTTGTTCTAATACTATTGGGC |
| **pNPTS CN-32 Δ*pilB*** | TR320 PilB KO1 | GAATTCGTGGATCCAGATATGTATAAGCTGGAGATAAATATGAAAGG |
| | TR321 PilB KO2 | TCGTCACCCGACCAGTGGTTGGCATAGATTCTTAA |
| | TR322 PilB KO3 | AACCACTGGTCGGGTGACGAGTTTTTAACAGC |
| | TR323 PilB KO4 | CAAGCTTCTCTGCAGGATCTTTTGGGCTCAATCTTCTTTGG |
| **pNPTS CN-32 Δ*mshE*** | AP241 EcoRV 0563 up fw | GAATTCGTGGATCCAGATGCTTACGCCAAGCCAGCTC |
| | AP242 OL_0563_up_rv | CCTCAACGCCCATCTTTAATCTGGGTTTCATTGGC |
| | AP243 OL_0563_down_fw | ATTAAAGATGGGCGTTGAGGCGTAATTATGC |
| | AP244 EcoRV_0563_down_rv | CAAGCTTCTCTGCAGGATCAAGGCAAATCGGCACCAAAG |
| **pNPTS CN-32 *mshA* S68C** | TR353 | GCGAATTCGTGGATCCAGATAAATGTAACCGACGACGCACAG |
| | TR357 | ATACATCCTTACACTCCACACCCTGAATAGCCG |
| | TR358 | GGGTGTGGAGTGTAAGGATGTATCTAGCATTATTATCGATG |

| | | |
|---|---|---|
| | TR356 | GCCAAGCTTCTCTGCAGGATGCTAGGC AGGCCTTTTCTAGTA |
| **pNPTS CN-32 *lapA*-GS-3x-FLAG** | VK239 EcoRV OL up 3591 fw 2 | GCGAATTCGTGGATCCAGATGGTGGTA GCCACAACGATGC |
| | VK240 up 3591 OL FLAG rv | AATATCATGATCTTTATAATCGCCATCAT GATCTTTATAATCACTGCCAGGGATCAT AGTGCCATTGTTATGAG |
| | VK241 OL FLAG 3591 down fw | GGCGATTATAAAGATCATGATATTGATT ATAAAGATGATGATGATAAATAAATAAAA TCGTTTTGATGGCCTATAGAAATATAGG |
| | VK233 EcoRV 3591-down rv | GCCAAGCTTCTCTGCAGGATGGCTTCTA GTGACTCAATATTGAGTGTC |
| **pNPTS CN-32 *lapB*-GS-3x-FLAG** | VK222 EcoRV OL up 3592 fw | GCGAATTCGTGGATCCAGATCCCTAGC GATCTACGCCG |
| | VK223 up 3592 OL FLAG rv | AATATCATGATCTTTATAATCGCCATCAT GATCTTTATAATCACTGCCTTTTTTACTG CCCCCATTGAACAG |
| | VK224 OL FLAG 3592 down fw | GATTATAAAGATCATGATATTGATTATAA AGATGATGATGATAAATAGGTTCAATGG GGGCAGTAAAAAATG |
| | VK225 EcoRV 3592 down rv | GCCAAGCTTCTCTGCAGGATCTCCGCC GCCACTATACTATC |
| **pNPTS CN-32 *pdeB-gfp* K527S** | TR564 pdeb OL plas fw | GCCAAGCTTCTCTGCAGGATGCAAGGC AATATGGATCCATCC |
| | TR568 K to S rv | TGATCTGGCTTAACAACTGCACCACAGA TAAAGC |
| | TR579 K to S fw | GCAGTTGTTAAGCCAGATCAGTGCTCAA GTCTCATTACAAG |
| | TR567 pdeB OL plas rv | GCGAATTCGTGGATCCAGATGCCAAAG ACGCGACTACAACTA |
| **pNPTS CN-32 *pdeB-gfp* K527S Q528S** | TR564 pdeb OL plas fw | GCCAAGCTTCTCTGCAGGATGCAAGGC AATATGGATCCATCC |
| | TR565 | TGATGCTGCTTAACAACTGCACCACAGA TAAAGC |
| | TR566 | GCAGTTGTTAAGCAGCATCAGTGCTCAA GTCTCATTACAAG |
| | TR567 pdeB OL plas rv | GCGAATTCGTGGATCCAGATGCCAAAG ACGCGACTACAACTA |
| **pNPTS CN-32 *pdeB-gfp* K527D** | TR244 ol pdeB up | GCCAAGCTTCTCTGCAGGATGCAAGGC AATATGGATCCATCC |
| | TR399 | TGATCTGGTCTAACAACTGCACCACAGA TAAAGC |
| | TR400 | GGTGCAGTTGTTAGACCAGATCAGTGC TCAAGTCTCATTA |
| | TR247 rv PdeB down | GCGAATTCGTGGATCCAGATGCCAAAG ACGCGACTACAACTA |
| **pNPTS CN-32 *pdeB-gfp* K527D Q528S** | TR244 ol pdeB up | GCCAAGCTTCTCTGCAGGATGCAAGGC AATATGGATCCATCC |
| | TR344 KQ to ES rv | TGATCGAGTCTAACAACTGCACCACAGA TAAAGCACTGCGAT |
| | TR345 KQ to ES fw | GCAGTTGTTAGACTCGATCAGTGCTCAA GTCTCATTACAAG |
| | TR247 rv PdeB down | GCGAATTCGTGGATCCAGATGCCAAAG ACGCGACTACAACTA |
| **pNPTS CN-32 *pdeB-gfp* Q524A K527D Q528A** | TR244 ol pdeB up | GCCAAGCTTCTCTGCAGGATGCAAGGC AATATGGATCCATCC |
| | TR281 | TGCGTCTAACAATGCCACCACAGATAAA GCACTGCGA |
| | TR282 | GCATTGTTAGACGCAATCAGTGCTCAAG TCTCATTACAAG |

| | | |
|---|---|---|
| | TR247 rv PdeB down | GCGAATTCGTGGATCCAGATGCCAAAG ACGCGACTACAACTA |
| **pNPTS CN-32 *pdeB-gfp* G497A** | TR564 pdeb OL plas fw | GCCAAGCTTCTCTGCAGGATGCAAGGC AATATGGATCCATCC |
| | TR570 G rv | CCTGTGGAGCAAGGCAGGCTTGCATCA CTTTAG |
| | TR571 G fw | AGCCTGCCTTGCTCCACAGGAGTTATTG GGGCGGATTGGTGG |
| | TR567 pdeB OL plas rv | GCGAATTCGTGGATCCAGATGCCAAAG ACGCGACTACAACTA |
| **pNPTS CN-32 *pdeB-gfp* Q499S** | TR564 pdeb OL plas fw | GCCAAGCTTCTCTGCAGGATGCAAGGC AATATGGATCCATCC |
| | TR572 Q rxxd rv | CGCTTGGACCAAGGCAGGCTTGCATCA CTTTAG |
| | TR573 Q rxxd rv | AGCCTGCCTTGGTCCAAGCGAGTTATT GGGGCGGATTGGTGG |
| | TR567 pdeB OL plas rv | GCGAATTCGTGGATCCAGATGCCAAAG ACGCGACTACAACTA |
| **pNPTS CN-32 *pdeB-gfp* E500S** | TR564 pdeb OL plas fw | GCCAAGCTTCTCTGCAGGATGCAAGGC AATATGGATCCATCC |
| | TR574 E rxxd rv | TCTGTGGACCAAGGCAGGCTTGCATCA CTTTAG |
| | TR575 E rxxd rv | AGCCTGCCTTGGTCCACAGAGCTTATTG GGGCGGATTGGTGG |
| | TR567 pdeB OL plas rv | GCGAATTCGTGGATCCAGATGCCAAAG ACGCGACTACAACTA |
| **pNPTS CN-32 *pdeB-gfp* Q524S Q528S** | TR244 ol pdeB up | GCCAAGCTTCTCTGCAGGATGCAAGGC AATATGGATCCATCC |
| | TR347 SLLKS rv | TTAACAAACTCACCACAGATAAAGCACT GCGAT |
| | TR348 SLLKS fw | ATCTGTGGTGAGTTTGTTAAAGTCGATC AGTGCTCAAGTCTCATTACAAG |
| | TR247 rv PdeB down | GCGAATTCGTGGATCCAGATGCCAAAG ACGCGACTACAACTA |
| **pNPTS CN-32 *pdeB-gfp* K527E Q528S** | TR564 pdeb OL plas fw | GCCAAGCTTCTCTGCAGGATGCAAGGC AATATGGATCCATCC |
| | TR395* ES rv | TGATCGATTCTAACAACTGCACCACAGA TAAAGC |
| | TR396* ES fw | GCAGTTGTTAGAATCGATCAGTGCTCAA GTCTCATTACAAG |
| | TR567 pdeB OL plas rv | GCGAATTCGTGGATCCAGATGCCAAAG ACGCGACTACAACTA |
| **pNPTS CN-32 *pdeB-gfp* K490D Q493A** | TR244 ol pdeB up | GCCAAGCTTCTCTGCAGGATGCAAGGC AATATGGATCCATCC |
| | TR279 | TGCCATCACGTCAGCAACCATGGCCAA CATGC |
| | TR280 | GACGTGATGGCAGCCTGCCTTGGTCCA CAG |
| | TR247 rv PdeB down | GCGAATTCGTGGATCCAGATGCCAAAG ACGCGACTACAACTA |
| **pNPTS CN-32 *pdeB-gfp* R557G A558G P559G Y560G** | TR244 ol pdeB up | GCCAAGCTTCTCTGCAGGATGCAAGGC AATATGGATCCATCC |
| | TR252 | GCCACCCCCTCCACCAAAGGCGACACC GATACTT |
| | TR253 | GGAGGGGGTGGCATCAATGCCCAAGAG TTGTTGAA |
| | TR247 rv PdeB down | GCGAATTCGTGGATCCAGATGCCAAAG ACGCGACTACAACTA |
| **pNPTS CN-32 *pdeB-gfp* V522G V523G Q524G** | TR244 ol pdeB up | GCCAAGCTTCTCTGCAGGATGCAAGGC AATATGGATCCATCC |

| | | |
|---|---|---|
| | TR248 | AACCCCCTCCAGATAAAGCACTGCGATTACAAATCA |
| | TR249 | GGAGGGGGTTTGTTAAAGCAGATCAGTGCTCAAG |
| | TR247 rv PdeB down | GCGAATTCGTGGATCCAGATGCCAAAGACGCGACTACAACTA |
| **pNPTS CN-32 *pdeB-gfp* K490G V491G M492G Q593G** | TR244 ol pdeB up | GCCAAGCTTCTCTGCAGGATGCAAGGCAATATGGATCCATCC |
| | TR279 | TGCCATCACGTCAGCAACCATGGCCAACATGC |
| | TR280 | GACGTGATGGCAGCCTGCCTTGGTCCACAG |
| | TR247 rv PdeB down | GCGAATTCGTGGATCCAGATGCCAAAGACGCGACTACAACTA |
| **pNPTS CN-32 *pdeB-gfp* K578S** | TR564 pdeb OL plas fw | GCCAAGCTTCTCTGCAGGATGCAAGGCAATATGGATCCATCC |
| | TR600 cn32 Ksalt to S rv | CGCCCCCTTCGCGCTAGCAGCAAGACAGGCAATATCAG |
| | TR601 cn32 Ksalt to S fw | GCCTGTCTTGCTGCTAGCGCGAAGGGGGCGAATCAAAT |
| | TR567 pdeB OL plas rv | GCGAATTCGTGGATCCAGATGCCAAAGACGCGACTACAACTA |
| **pNPTS CN-32 *pdeB-gfp* K580S** | TR564 pdeb OL plas fw | GCCAAGCTTCTCTGCAGGATGCAAGGCAATATGGATCCATCC |
| | TR602 cn32 Kc to S rv | TTGATTCGCCCCGCTCGCTTTAGCAGCAAGACAGG |
| | TR603 cn32 Kc to S fw | CTTGCTGCTAAAGCGAGCGGGGCGAATCAAATCCATATTTATG |
| | TR567 pdeB OL plas rv | GCGAATTCGTGGATCCAGATGCCAAAGACGCGACTACAACTA |
| **pNPTS CN-32 *pdeB* G497A** | TR564 pdeb OL plas fw | GCCAAGCTTCTCTGCAGGATGCAAGGCAATATGGATCCATCC |
| | TR570 G rv | CCTGTGGAGCAAGGCAGGCTTGCATCACTTTAG |
| | TR571 G fw | AGCCTGCCTTGCTCCACAGGAGTTATTGGGGCGGATTGGTGG |
| | TR567 pdeB OL plas rv | GCGAATTCGTGGATCCAGATGCCAAAGACGCGACTACAACTA |
| **pNPTS CN-32 *pdeB* K578S** | TR564 pdeb OL plas fw | GCCAAGCTTCTCTGCAGGATGCAAGGCAATATGGATCCATCC |
| | TR600 cn32 Ksalt to S rv | CGCCCCCTTCGCGCTAGCAGCAAGACAGGCAATATCAG |
| | TR601 cn32 Ksalt to S fw | GCCTGTCTTGCTGCTAGCGCGAAGGGGGCGAATCAAAT |
| | TR567 pdeB OL plas rv | GCGAATTCGTGGATCCAGATGCCAAAGACGCGACTACAACTA |
| **pNPTS CN-32 *pdeB* K580S** | TR564 pdeb OL plas fw | GCCAAGCTTCTCTGCAGGATGCAAGGCAATATGGATCCATCC |
| | TR602 cn32 Kc to S rv | TTGATTCGCCCCGCTCGCTTTAGCAGCAAGACAGG |
| | TR603 cn32 Kc to S fw | CTTGCTGCTAAAGCGAGCGGGGCGAATCAAATCCATATTTATG |
| | TR567 pdeB OL plas rv | GCGAATTCGTGGATCCAGATGCCAAAGACGCGACTACAACTA |
| **pNPTS CN-32 *pdeB-mvenus*** | TR244 ol pdeB up | GCCAAGCTTCTCTGCAGGATGCAAGGCAATATGGATCCATCC |
| | TR456 PdeB-Venus up rv | CTCGCCCTTGCTCACTGCGCGTTGTGCTAAACCCATCTCA |
| | TR457 Venus OL PdeB fw | TTAGCACAACGCGCAGTGAGCAAGGGCGAGGAGCTGTTCA |

| | | |
|---|---|---|
| | TR458 Venus OL PdeB fw | AGCGCAAATTCATCACTTGTACAGCTCG TCCATGCCGAGA |
| | TR459 PdeB-Venus dn fw | GACGAGCTGTACAAGTGATGAATTTGC GCTTTTAGTCCGA |
| | TR247 rv PdeB down | GCGAATTCGTGGATCCAGATGCCAAAG ACGCGACTACAACTA |
| **pNPTS CN-32 *pdeB-venus* D508A E509A** | TR244 ol pdeB up | GCCAAGCTTCTCTGCAGGATGCAAGGC AATATGGATCCATCC |
| | TR456 PdeB-Venus up rv | CTCGCCCTTGCTCACTGCGCGTTGTGC TAAACCCATCTCA |
| | TR457 Venus OL PdeB fw | TTAGCACAACGCGCAGTGAGCAAGGGC GAGGAGCTGTTCA |
| | TR458 Venus OL PdeB fw | AGCGCAAATTCATCACTTGTACAGCTCG TCCATGCCGAGA |
| | TR459 PdeB-Venus dn fw | GACGAGCTGTACAAGTGATGAATTTGC GCTTTTAGTCCGA |
| | TR247 rv PdeB down | GCGAATTCGTGGATCCAGATGCCAAAG ACGCGACTACAACTA |
| **pNPTS CN-32 *pdeB-venus* E637A** | TR244 ol pdeB up | GCCAAGCTTCTCTGCAGGATGCAAGGC AATATGGATCCATCC |
| | TR456 PdeB-Venus up rv | CTCGCCCTTGCTCACTGCGCGTTGTGC TAAACCCATCTCA |
| | TR457 Venus OL PdeB fw | TTAGCACAACGCGCAGTGAGCAAGGGC GAGGAGCTGTTCA |
| | TR458 Venus OL PdeB fw | AGCGCAAATTCATCACTTGTACAGCTCG TCCATGCCGAGA |
| | TR459 PdeB-Venus dn fw | GACGAGCTGTACAAGTGATGAATTTGC GCTTTTAGTCCGA |
| | TR247 rv PdeB down | GCGAATTCGTGGATCCAGATGCCAAAG ACGCGACTACAACTA |
| **pNPTS MR-1 Δ*pdeB*** | TR582 SO pdeB KO cterm500 | GCCAAGCTTCTCTGCAGGATGCCAAGC CATAATCTTATGCTTTAGG |
| | TR583 SO pdeB KO start | GTTGTGCTAAGTTGCCTATGCGCATCTT TTACC |
| | TR584 SO pdeB KO stop | CATAGGCAACTTAGCACAACGCGCATA GGG |
| | TR581 SO pdeB nterm500 | GCGAATTCGTGGATCCAGATTAACAGCA TGTTTAGACGCCGC |
| **pNPTS MR-1 *pdeB-gfp*** | TR576 SO up fw | GCCAAGCTTCTCTGCAGGATCCGCAGC AGAGCGTTTTAAGC |
| | TR577 SO pdeb nterm rv | TGCTGCTGCCTGCGCGTTGTGCTAAGC GC |
| | TR578 SO pdeb-gfp fw | ACAACGCGCAGGCAGCAGCAAAGGAGA AGAACTTTTC |
| | TR579 SO pdeb-gfp rv | CAATCCCCTAGGATCCTTTGTAGAGCTC ATCC |
| | TR580 SO pdeb nterm fw | CAAAGGATCCTAGGGGATTGCGCTTTTA AGGTG |
| | TR581 SO pdeB nterm500 | GCGAATTCGTGGATCCAGATTAACAGCA TGTTTAGACGCCGC |
| **pNPTS MR-1 *pdeB-gfp* K524E Q525S** | TR582 SO pdeB KO cterm500 | GCCAAGCTTCTCTGCAGGATGCCAAGC CATAATCTTATGCTTTAGG |
| | TR593 SO KQ to ES rv | TAATGCTTTCCAGCAACTGCGCCACAGC TAA |
| | TR594 SO KQ to ES fw | GCAGTTGCTGGAAAGCATTATTGCTCAA GTATCGCTGCAAGTG |
| | TR581 SO pdeB nterm500 | GCGAATTCGTGGATCCAGATTAACAGCA TGTTTAGACGCCGC |
| **pNPTS MR-1 *pdeB-gfp* G494A** | TR582 SO pdeB KO cterm500 | GCCAAGCTTCTCTGCAGGATGCCAAGC CATAATCTTATGCTTTAGG |

| | | |
|---|---|---|
| | TR544 G494A fw | ATTCCTGTGGCGCAAGACATGACTGAATCGCCCTAG |
| | TR555 G494A rv | ATGTCTTGCGCCACAGGAATTATTAGCCCGCA |
| | TR581 SO pdeB nterm500 | GCGAATTCGTGGATCCAGATTAACAGCATGTTTAGACGCCGC |
| **pNPTS MR-1 *pdeB-gfp* K575S** | TR582 SO pdeB KO cterm500 | GCCAAGCTTCTCTGCAGGATGCCAAGCCATAATCTTATGCTTTAGG |
| | TR598 aSak rv | GGTGCCCTTGGCACTAGCGGCAATACAGGCGATATCT |
| | TR599 aSak fw | GCCTGTATTGCCGCTAGTGCCAAGGGCACCAACCAAAT |
| | TR581 SO pdeB nterm500 | GCGAATTCGTGGATCCAGATTAACAGCATGTTTAGACGCCGC |
| **pNPTS MR-1 *pdeB-gfp* K577S** | TR582 SO pdeB KO cterm500 | GCCAAGCTTCTCTGCAGGATGCCAAGCCATAATCTTATGCTTTAGG |
| | TR596 akaS rv | TTGGTTGGTGCCACTGGCTTTAGCGGCAATACAGG |
| | TR597 akaS fw | ATTGCCGCTAAAGCCAGTGGCACCAACCAAATCCACATTTA |
| | TR581 SO pdeB nterm500 | GCGAATTCGTGGATCCAGATTAACAGCATGTTTAGACGCCGC |

### Additional References


1. V. L. Miller, J. J. Mekalanos, A novel suicide vector and its use in construction of insertion mutations: osmoregulation of outer membrane proteins and virulence determinants in *Vibrio cholerae* requires *toxR*. *J. Bacteriol.* **170**, 2575–2583 (1988).

2. J. K. Fredrickson, *et al.*, Biogenic iron mineralization accompanying the dissimilatory reduction of hydrous ferric oxide by a groundwater bacterium. *Geochim. Cosmochim. Acta* **62**, 3239–3257 (1998).

3. S. Bubendorfer, *et al.*, Specificity of motor components in the dual flagellar system of *Shewanella putrefaciens* CN-32. *Mol. Microbiol.* **83**, 335–350 (2012).

4. F. M. Rossmann, *et al.*, The GGDEF Domain of the phosphodiesterase PdeB in *Shewanella putrefaciens* mediates recruitment by the polar landmark protein HubP. *J. Bacteriol.* **201** (2019).

5. K. Venkateswaran, *et al.*, Polyphasic taxonomy of the genus *Shewanella* and description of *Shewanella oneidensis* sp. nov. *Int. J. Syst. Bacteriol.* **49 Pt 2**, 705–724 (1999).

6. J. Lassak, A.-L. Henche, L. Binnenkade, K. M. Thormann, ArcS, the cognate sensor kinase in an atypical Arc system of *Shewanella oneidensis* MR-1. *Appl. Environ. Microbiol.* **76**, 3263–3274 (2010).

7. F. Rossmann, *et al.*, The role of FlhF and HubP as polar landmark proteins in *Shewanella putrefaciens* CN-32. *Mol. Microbiol.* **98**, 727–742 (2015).

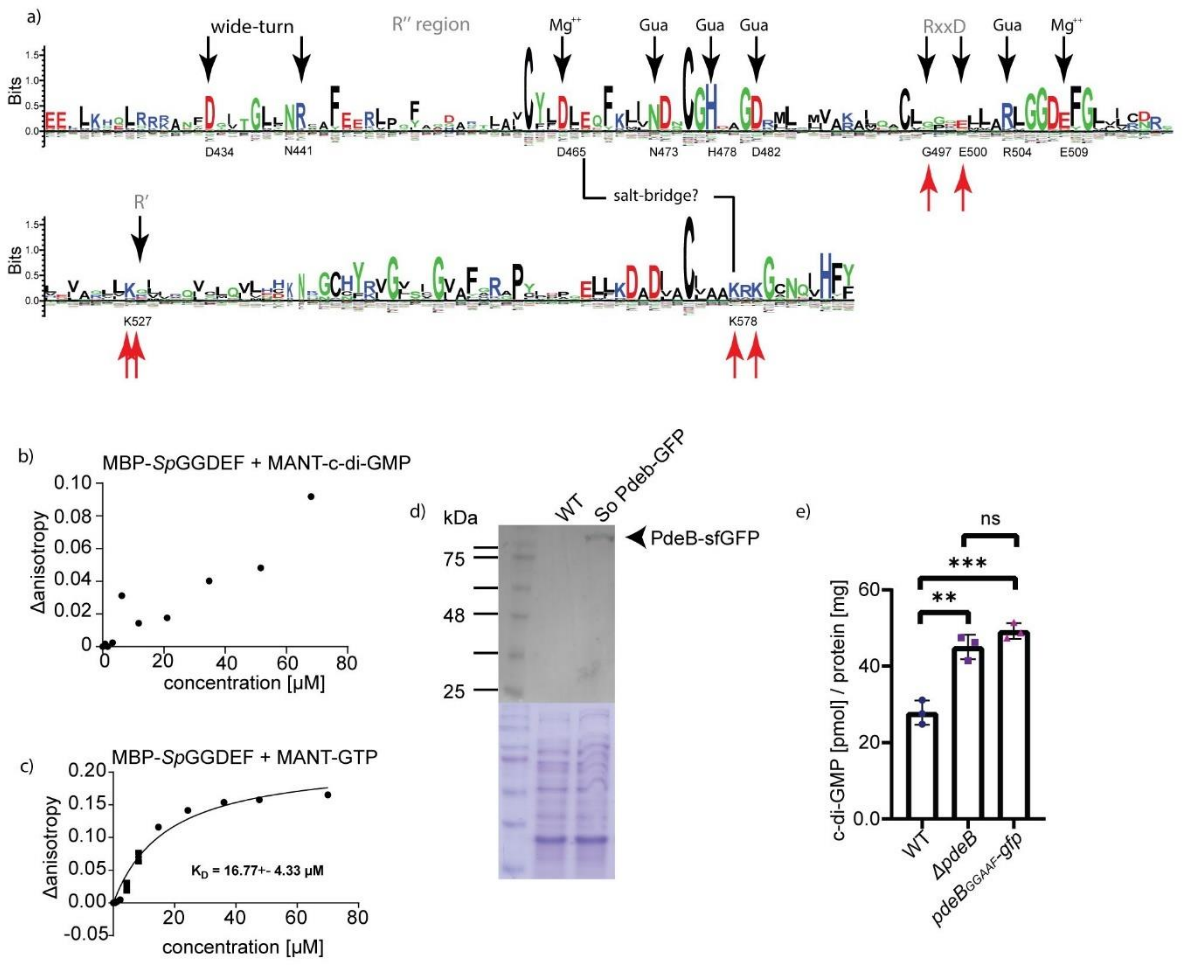


**Supplemental Figure 1. a)** Position-based weight map of the GGDEF domain of 50 PdeB homologues from different *Shewanella* species. Characteristic features of GGDEF domains are marked with black arrows and degenerated or missing motifs are indicated in gray. Residues that are important for the GGDEF$_{PdeB}$-FimVc$_{HubP}$ interaction are highlighted by red arrows. **b)** The MANT-c-di-GMP binding of the GGDEF domain of *So*PdeB was tested by fluorescence anisotropy assays. No binding curve was observed, only unspecific binding occurred at unphysiological high ligand concentrations. **c)** Binding of MANT-GTP to the GGDEF domain of *So*PdeB was tested using fluorescence anisotropy assays. The assay confirmed binding with a $K_D$ value around 15 µM. **d)** The stability and expression of genomic *So*PdeB-sfGFP fusions was verified by immunoblot analysis. **e)** The effect of GTP binding to GGDEF$_{PdeB}$ on the PDE activity of PdeB was tested introducing mutations the GGDEF motif. The cellular c-di-GMP was then extracted and quantified by mass spectroscopy. The single point mutation into the GGDEF motif leads to the same increase in cellular c-di-GMP concentrations as deletion of *pdeB*.

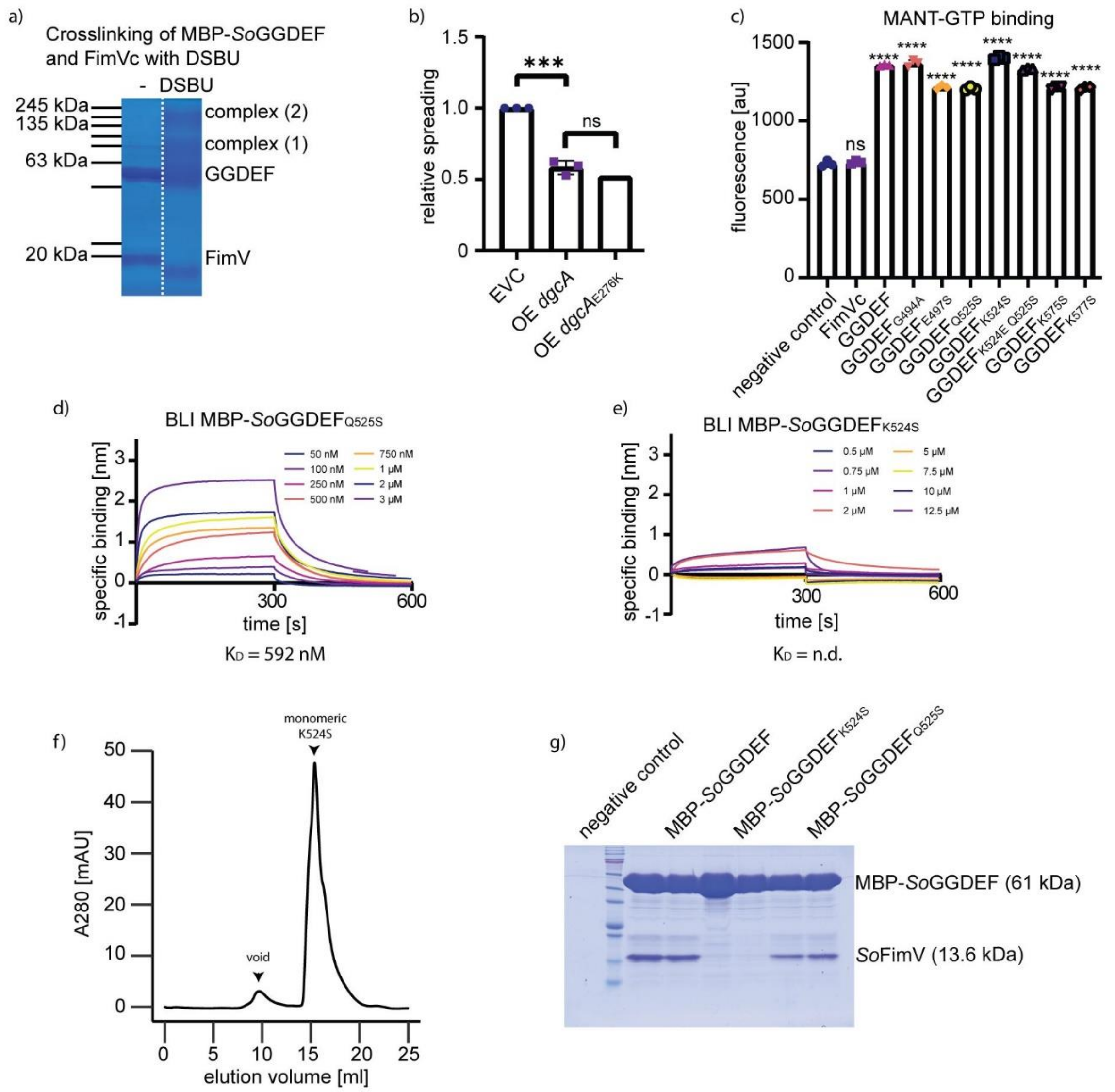


**Supplemental Figure 2: Structural and sequential features of PdeB$_{GGDEF}$ and HubP$_{FimV}$. a)** The crosslinking of *So*GGDEF with *So*HubP was verified using SDS-PAGE. **b)** Mutating the conserved glutamic acid in the R'' site does not inhibit the DGC activity of DgcA as shown with soft-agar motility assays. **c)** Functionality of MBP-*So*GGDEF proteins used for BLI was shown by MANT-GTP binding assays. All mutated versions are able to bind MANT-GTP, as indicated by the increased fluorescence. **d-e)** BLI assays for GGDEF mutants with substitutions in the R` I-site show decreased affinity to FimV compared to the wild type. The purified MBP-GGDEF$_{K524S}$ showed aggregation upon production and unspecific binding in BLI assays and was therefore not suitable for determination of exact KD values. **f)** The stability was therefore tested by storing the protein for three days at 4°C and further analysis using SEC. The sample remained mostly in monomeric form. **g)** Interaction was tested with pulldown assays were the wild-type version and MBP-GGDEF$_{Q525S}$ served as controls. No binding was observed when K524 was mutated.

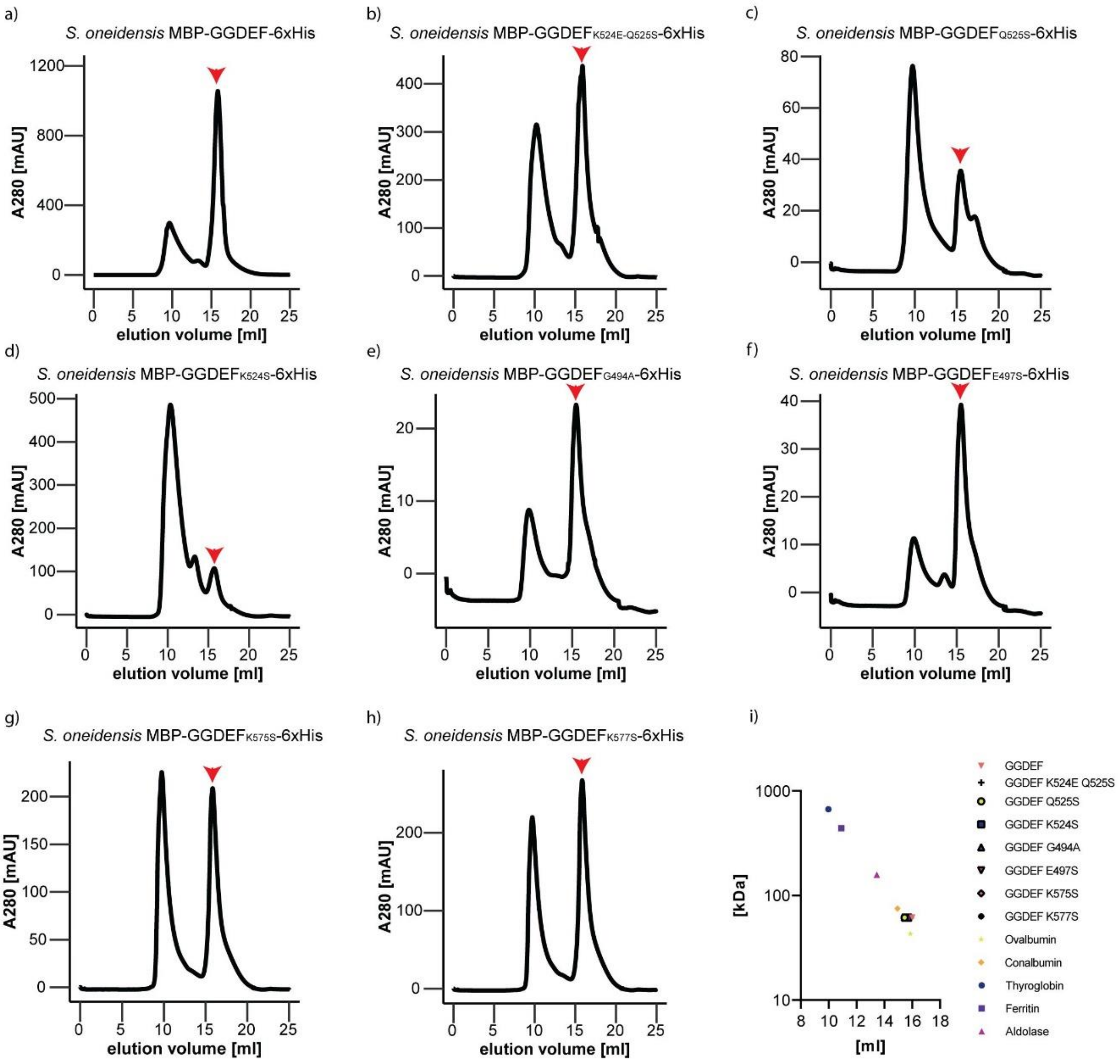


**Supplemental Figure 3: SEC of GGDEF domains. a-h)** The mutated $GGDEF_{PdeB}$ domains of *S. oneidensis* were purified as MBP-fusion proteins. The chromatograms are shown in a-h, were the wild type version (a) serves as positive control. The elution volume in ml is plotted against the absorbance at 280 nm. Red arrows indicate the peak for the monomeric proteins of interest. **i)** The elution volume of the proteins of interests was plotted against the molecular weight in kDa, together with globular proteins included in the high molecular weight calibration kit (GE healthcare). All GGDEF proteins elute at roughly the same elution volume, indicating that the structure is not altered.

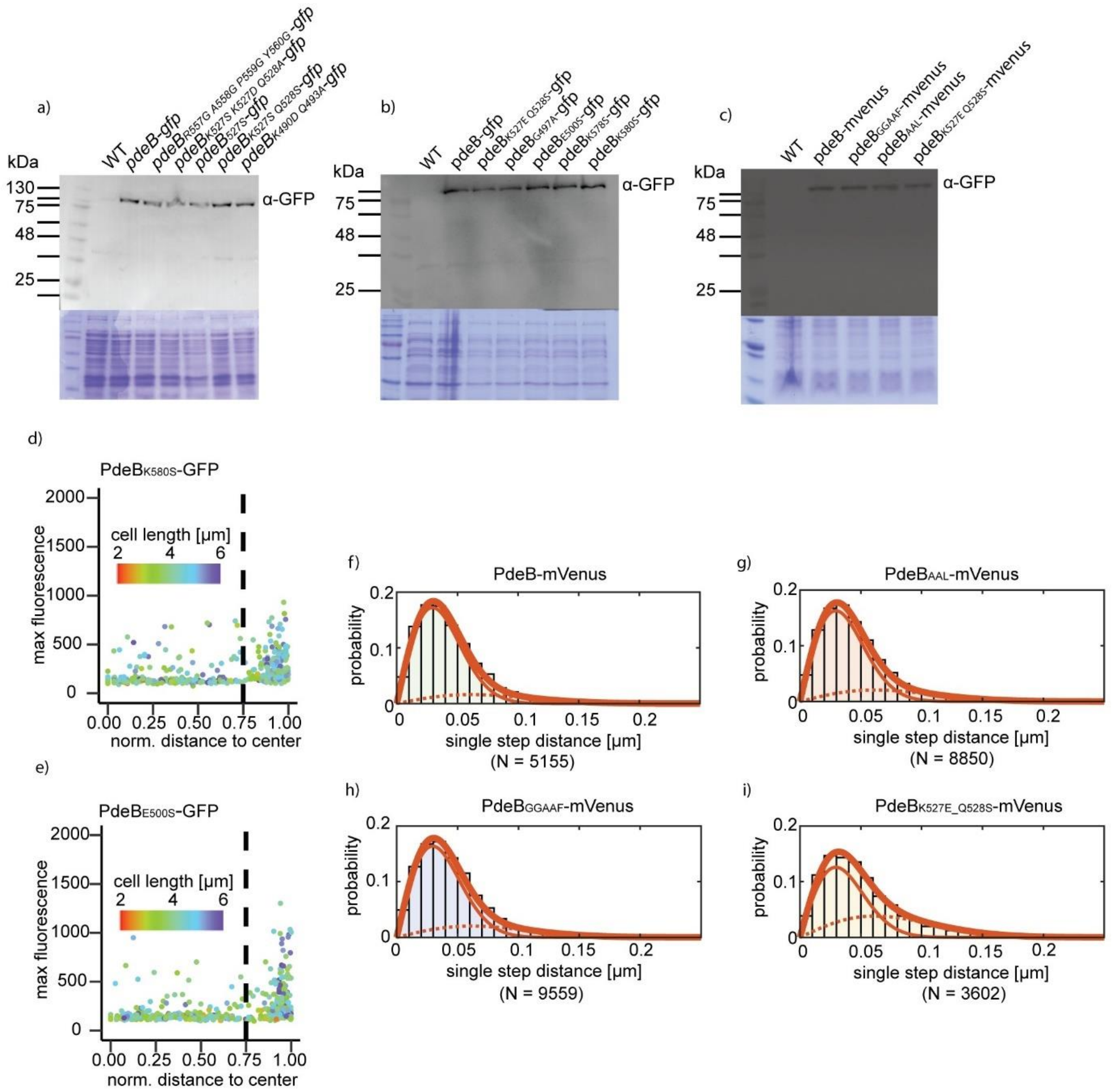


**Supplemental Figure 4. a-b)** Stability and expression of *Sp*PdeB-sfGFP mutants were verified using Western blot analysis. **c)** Stability of *Sp*PdeB-mVenus used for single molecule microscopy was shown by Western blot analysis. **d-e)** Scatterplots of fluorescence microscopy of mutants with reduced PdeB-sfGFP localization. Both mutations reduce the polar localization of PdeB-sfGFP, likely due to reduced affinity to HubP. **f-i)** Fits for the jump distance analyses used for the data of the bubble blot. Solid thin line, Rayleigh fit for the slow population; dotted line, fast population; thick solid line, combination of both fits. $R^2$ of values all fits were higher than 0.999. For all mVenus fusions, a two population-fit was the best.

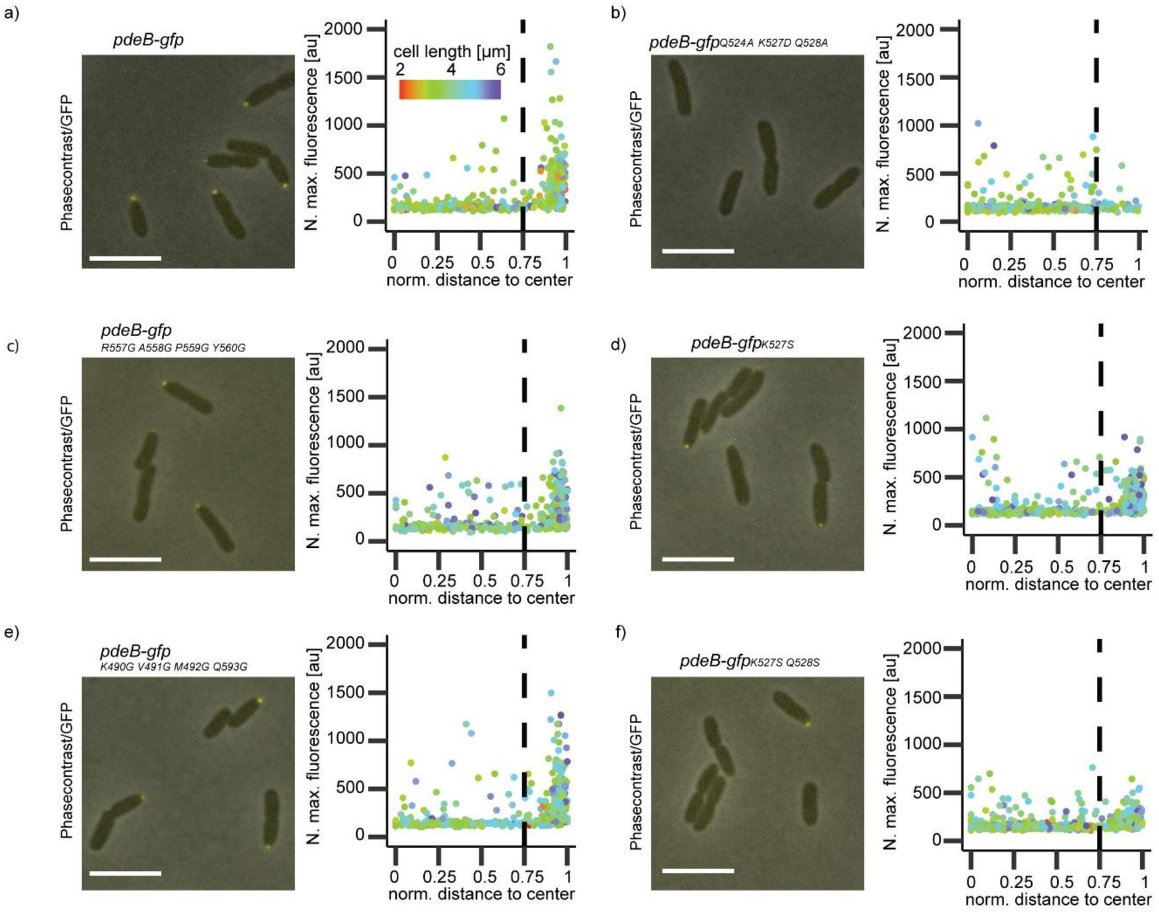


**Supplemental Figure 5. Screening for residues involved in the polar localization of PdeB. a-f)** Residues at different regions in the GGDEF domain of *Sp*PdeB-sfGFP were genomically mutated and localization behavior was observed using fluorescence microscopy. The localization behaviors of mutants are shown as scatter plots and wild-type *Sp*PdeB-sfGFP serves as control. Mutating residues of the R` site (b, d, f) leads to reduced polar localization, while the other two regions only had minor effects.

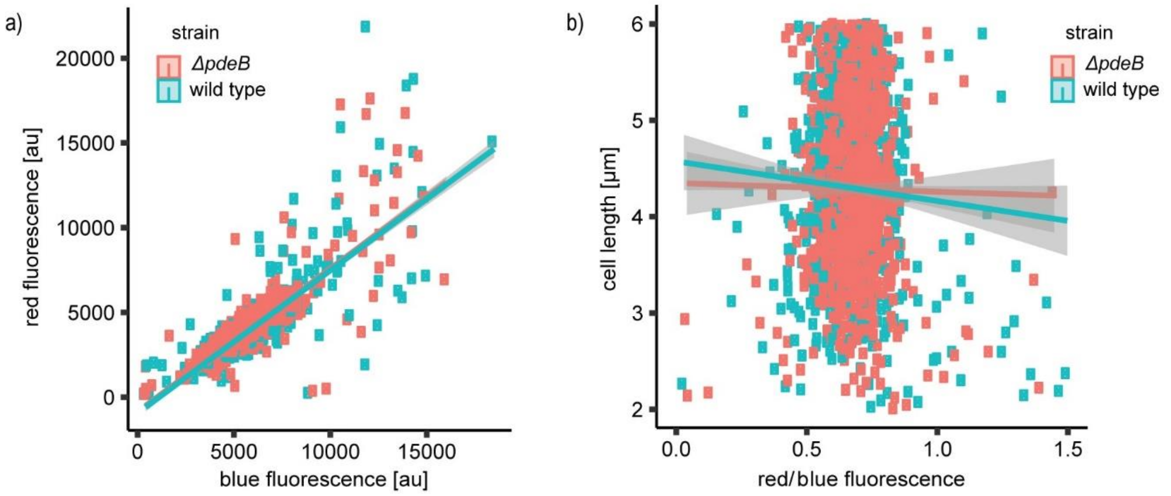


**Supplemental Figure 6: c-di-GMP single cell reporter controls. a)** The functionality of the fluorescence-based c-di-GMP reporter was tested for *S. putrefaciens* by plotting the blue against the red fluorescence and testing for linear correlation in presence and absence of *pdeB*. **b)** A correlation of cell length with the c-di-GMP level was tested by plotting the quotient of the red fluorescence divided by the blue fluorescence against the cell length. However, no correlation was found.

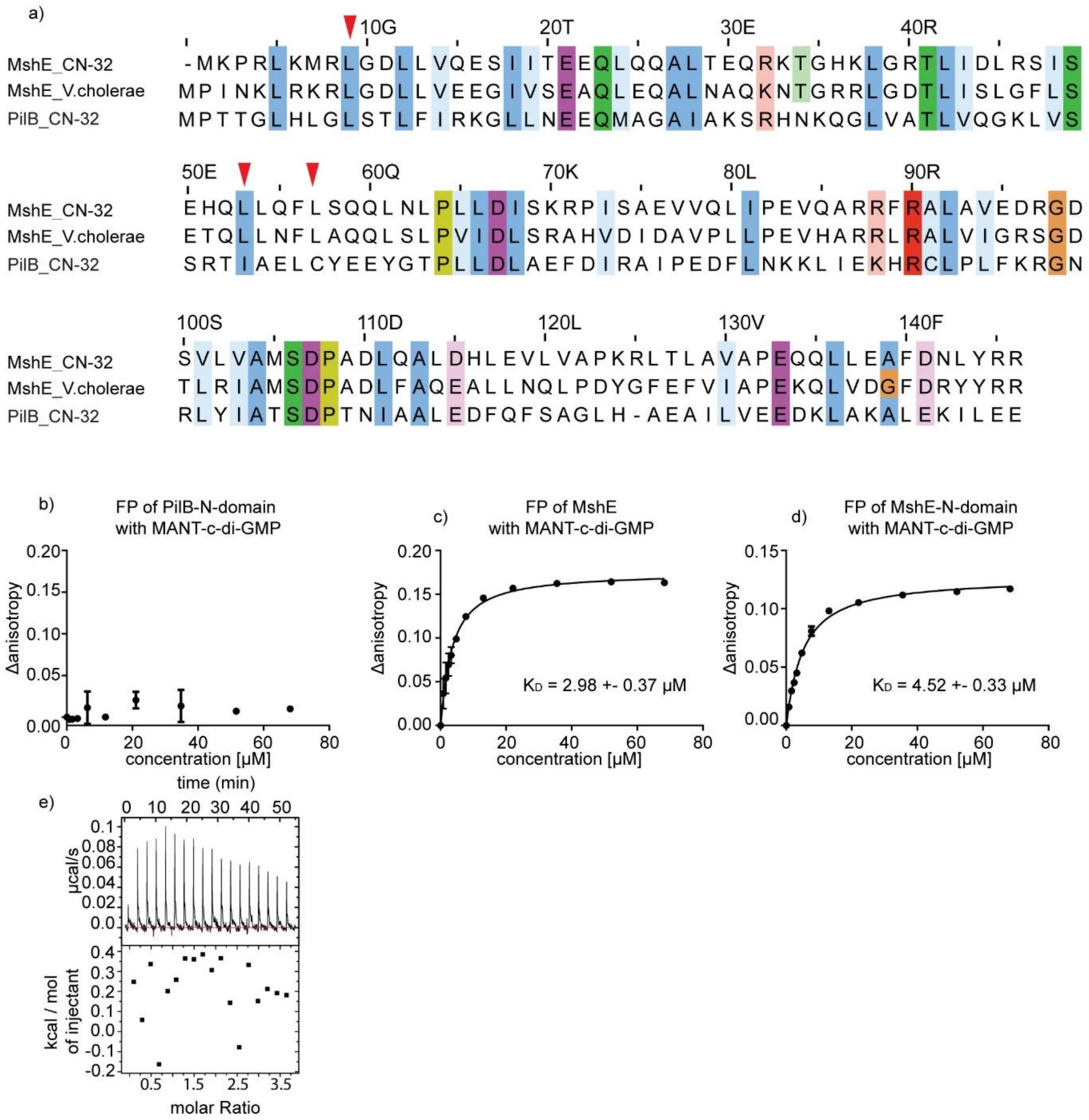


**Supplemental Figure 7. a)** Alignment of *Pa*MshE with *Sp*MshE and *Sp*PilB. Leucines that are involved in c-di-GMP binding are marked with red arrows. **b-d)** The MANT-c-di-GMP binding of the N-terminal domain of PilB (b), MshE (c) and the N-terminal domain of MshE (d) was tested by fluorescence anisotropy measurements. Both MshE versions are able to bind MANT-c-di-GMP with high affinity, while PilB does not. **e)** The non-binding of c-di-GMP to PilB was verified by ITC.

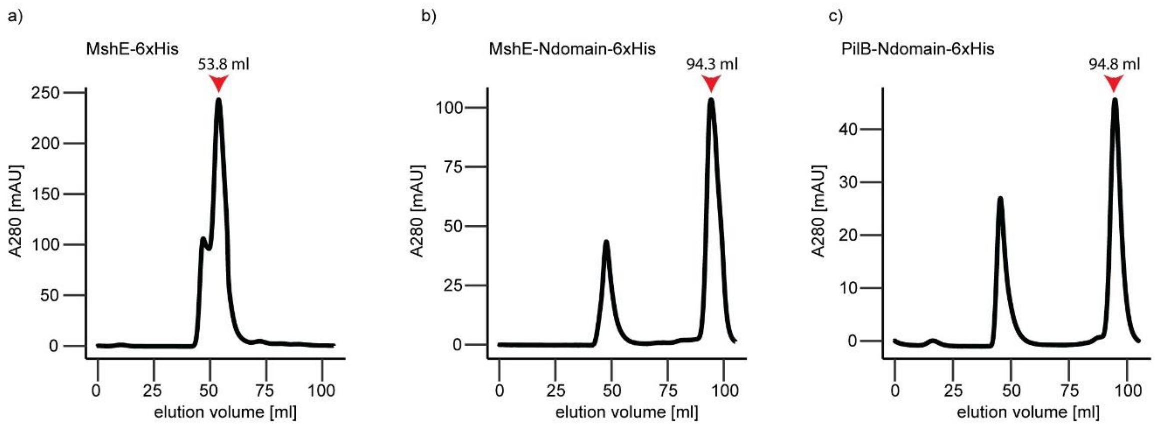


**Supplemental Figure 8: SEC of MShE and PilB. a-b)** Chromatograms of the size-exclusion chromatography of the extension ATPase MshE. Chromatograms show the elution volume against the absorbance at 280 nm. Peaks that contain the protein of interest are indicated by red arrows. The full length MshE protein (a) eluted at 53.8 ml, indicating an oligomeric state (penta- or hexameric), while the N-terminal domain eluted as monomer. **c)** The N-terminal domain of PilB eluted as monomer.

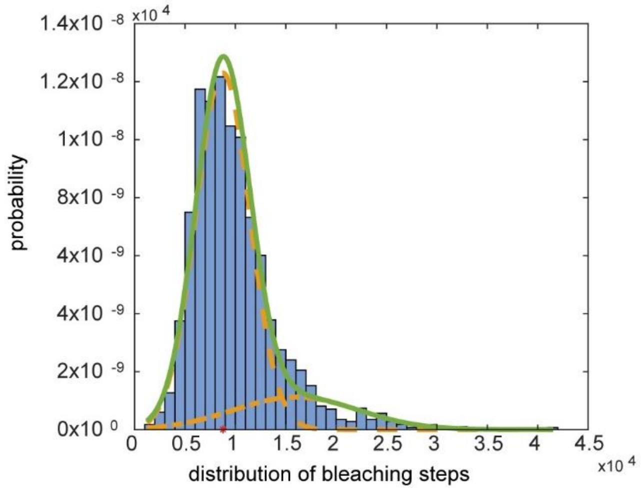


**Supplemental Figure 9: Fluorescence-based molecule quantification of PdeB-mVenus.** The distribution of bleaching steps within the movies is shown as histogram.
